# Dissecting origins of wiring specificity in dense reconstructions of neural tissue

**DOI:** 10.1101/2024.12.14.628490

**Authors:** Philipp Harth, Daniel Udvary, Jan Boelts, Jakob H. Macke, Daniel Baum, Hans-Christian Hege, Marcel Oberlaender

## Abstract

What are the origins of the highly specific wiring patterns that are formed by the neurons in the brain? To address this question, we introduce a method to predict the empirically observed wiring diagram – the connectome – at synaptic resolution based on dense electron-microscopic reconstructions of neural tissue. Our method generates the connectome based on the morphological properties of the neurons in combination with *synaptic specificity* models, whose parameters capture how neurons wire conditional on their subcellular, cellular and cell type properties. We employ simulation-based Bayesian inference to identify all values for these parameters that can generate the observed connectome. Finally, for each synapse in a dense reconstruction, our method provides quantitative measures to reveal which synaptic specificity models are necessary, sufficient and best-suited to generate it. The output of our method are experimentally testable predictions of wiring preferences from subcellular to cell type levels that could account for each synapse in dense reconstructions. We demonstrate our method on dense datasets from mouse primary visual and human temporal cortex. Strikingly, this demonstration shows that just three assumptions, with nearly the same synaptic specificity values, predict the connectivity in both datasets. Our method is openly accessible as a computational framework that includes a comprehensive workflow for the analysis of wiring specificity, and which provides users with full flexibility to define and test their hypotheses. Our method sets the stage to uncover the principles by which neural networks are organized, and to compare these principles across brain areas, species and time points.

## Introduction

The neurons in the brain are not randomly interconnected. Empirical evidence for this so-called ‘wiring specificity’^1^ has been observed across scales, from subcellular to network levels. For example, at the network scale, the connectivity patterns between three or more neurons form motifs that occur more or less frequently than in random networks with the same pairwise connectivity^2–5^. At the cellular scale, connection probabilities between pairs of neurons generally depend on their cell types^6^, inter-somatic distances^7^, and other cellular properties, such as the brain areas targeted by their respective long-range axons^8^. At the subcellular scale, neurons often form multi-synaptic connections with other neurons^9^, sometimes as clusters of synapses on the cell body (i.e., soma), same dendritic branch, or axon initial segment (AIS)^10–14^. How origins of such wiring specificity can be uncovered remains an open question.

Addressing this question is difficult because multiple different sources can be the origin for the highly specific wiring patterns between neurons. On one hand, synaptic connections are constantly formed, eliminated and replaced conditional on different neuronal properties^3, 15, 16^. These properties can reflect intrinsic factors of the pre- and postsynaptic neurons, such as their genetic cell types, extrinsic factors, such as their activity patterns, or combinations thereof^17–19^. On the other hand, the morphological properties of the neurons constrain which neuronal structures are sufficiently close to potentially form connections^20^. Moreover, beyond merely constraining connectivity, we previously showed that neuron morphology by itself is a major source for wiring specificity across all scales^5^. In essence, even if the molecular mechanisms that wire neurons conditional on their properties were absent, networks formed by real neuron morphologies will inevitably display non-random network motifs, cell type- and location-specific connectivity, as well as multi-synaptic clusters^5^. Dissecting the origins of wiring specificity hence faces two major challenges. First, it requires the ability to reconstruct the connectivity between complete neuron morphologies at synaptic (i.e., nanoscale) resolution. Second, it requires the ability to test which wiring rules – i.e., wiring conditional on which neuronal properties – account best for which of the observed connections.

Since its introduction by Denk & Horstmann in 2004^21^, dense electron-microscopic reconstruction has been a promising approach for addressing the first challenge^22^. Indeed, recent advances in automated tracing and proof-reading algorithms have started to yield data at nanoscale resolution that describes the connectivity between the neuronal structures of all neurons within large volumes – e.g. of a cortical ‘barrel’ column in mouse somatosensory cortex^23^, or of an entire fruit fly brain^24^. These breakthroughs in the field of dense reconstruction indicate that it is now the time to tackle the second challenge.

Indeed, first studies started to tackle this second challenge by applying computational models to dense reconstructions. For example, Schneider-Mizell et al., tested whether neuron morphology – modeled by proximity between pre- and postsynaptic structures – could account for inhibitory synapses from chandelier cells along AISs of pyramidal neurons^13^. When this proximity-based model was applied to a dense dataset from layer 2/3 (L2/3) of mouse primary visual cortex, it predicted synapse distributions that deviated from those observed empirically, suggesting that chandelier cell axons wire with AISs conditional on soma size and somatic inhibition of the respective pyramidal neurons^13^. Motta et al., examined whether the electrical activity of the pre- and postsynaptic neurons in the form of Hebbian synaptic plasticity – modelled by synaptic weight similarity – could account for multi-synaptic connections in a dense reconstruction from layer 4 (L4) of the mouse primary somatosensory cortex^12^. This synaptic weight-based model predicted upper bounds of 10 to 20% for the fraction of connections that could have undergone long-term potentiation or long-term depression^12^.

These examples illustrate that modelling wiring specificity in dense reconstructions has the potential to uncover wiring rules that provide insights into the principles by which a neural network is organized, and eventually into mechanisms by which it grows and develops^25^. However, these examples also illustrate the limits of current modelling approaches. First, proximity-based models – often referred to as “Peters’ rule”^26^ – are generally insufficient to assess how neuron morphology impacts connectivity^5^ (see also Discussion). Second, current models investigate one connectivity feature – e.g. distributions of inhibitory synapses onto AISs^13^ or occurrences of multi-synaptic connections^12^. Whether the model assumptions could also account for other connectivity features of the neurons at cellular and network scales remains unclear. Third, current models test only one assumption as the potential origin of wiring specificity – e.g. proximity^13^ or Hebbian plasticity^12^. It remains hence unclear whether other model assumptions account equally well, or even better, for the observed connectivity patterns. Moreover, the origins of the wiring specificity that are not accounted for by the tested assumption remain unclear. For example, Hebbian plasticity could not explain more than 80% of the multi-synaptic connections in the dense dataset from L4 of mouse somatosensory cortex^12^.

Here, we introduce a method that overcomes the present limitations for dissecting the origins of wiring specificity. We demonstrate our method by applying it to recently reported dense reconstructions from mouse primary visual cortex^27^ and human temporal cortex^14^. Our method is openly accessible via a computational framework that provides a comprehensive workflow for analyzing the origins of wiring specificity from subcellular to network scales, and which provides users with full flexibility to define and test their own hypotheses.

## Results

Our computational framework requires four inputs (**Fig. 1a**). The first input is the dense reconstruction. The second input is the connectome observed from this dense reconstruction. Together, these inputs provide the 3D locations of all neuronal structures and all synapses between them. The third input specifies which connectivity patterns the framework considers for analysis. Users can define these connectivity patterns at network, cellular or subcellular scales via an interface. Here, we consider the numbers of synapses from and to excitatory (EXC) and inhibitory (INH) neurons, motifs formed by three neurons, whether neurons connect on a soma, dendrite or AIS, and by how many synapses. The fourth input defines the assumptions tested by the framework. For this purpose, we provide an interface where users can define *synaptic specificity* parameters α_i_ ∈ [−1, +1]; i = 1 … *k*, which model wiring conditional on neuronal properties at population (P), cellular (C) and subcellular (S) level^17, 18, 28^. Here, we consider that neurons may exhibit negative or positive preferences for connecting to one another conditional on their cell type (α_*P*i_), cellular identity (α_*C*i_), subcellular targets (α_*S*i_), and combinations thereof (α_*PSC*i_). Thus, our framework enables users to define and test various biologically interpretable synaptic specificity assumptions across different connectivity patterns in dense reconstructions. This analysis is organized into a five-step workflow (see Materials and Methods for more details).

**Figure 1:**
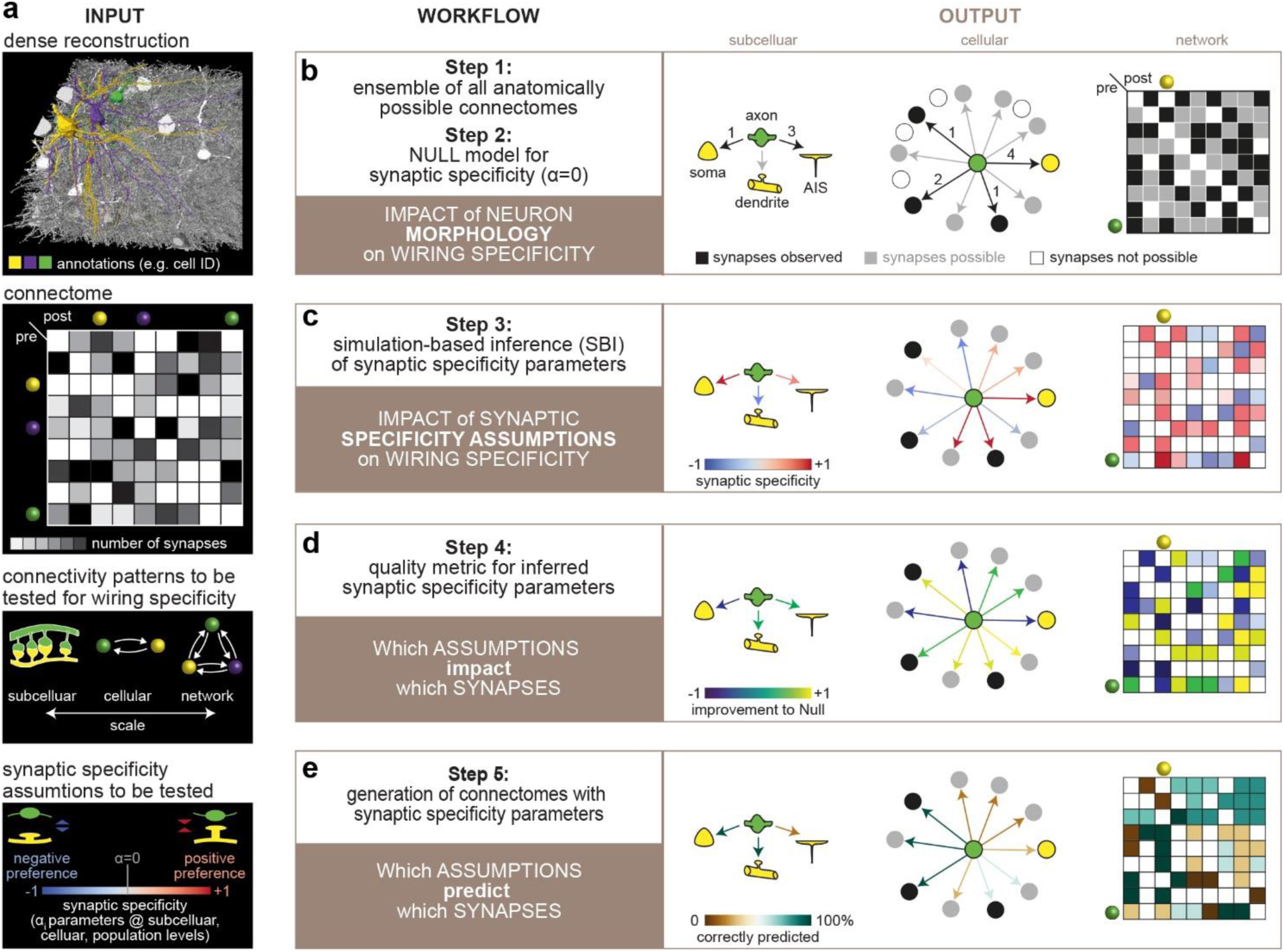
Schematic illustration of our method for dissecting origins of wiring specificity. **(a)** Our computational framework requires four inputs. From top to bottom: (1) a dense reconstruction of neural tissue, (2) the connectome obtained from this dense reconstruction, (3) a definition of the connectivity patterns that the framework should consider for analysis (i.e., users specify these patterns at network, cellular and subcellular scales via an interface), (4) a definition of the assumptions that the framework will test – i.e., users define *synaptic specificity* parameters α_i_ ∈ [−1, +1]; i = 1 … *k* that model wiring conditional on neuronal properties at population (P), cellular (C) or subcellular (S) levels. **(b-e)** Our analysis is structured into a five-step workflow. **(b)** Step 1 generates an *ensemble of connectomes*^5^ by modelling *possible synapses* as all combinations by which the pre- and postsynaptic sites in the data could connect to one another. For this purpose, users set a distance beyond which connections are not possible. Step 2 tests the null hypothesis that *“synaptic specificity is not the origin of the connectivity patterns*” – i.e., we set all α_i_ parameters to zero. The null hypothesis reveals the impact of neuron morphology on wiring specificity^5^. **(c)** Step 3 tests the impact of synaptic specificity assumptions on wiring specificity. For this purpose, we consider α_i_ parameters as free model parameters and use simulation-based Bayesian inference (SBI)^29^ to identify those α_i_ values that account best for the tested connectivity patterns. **(d)** Step 4 evaluates the impact of the synaptic specificity assumptions on each synapse in the data – i.e., we quantify via a *loss function* how the likelihood for each possible synapse to occur changes across different synaptic specificity assumptions. **(e)** Step 5 tests whether the inferred α_i_ values could account for the entire empirically observed connectome – i.e., we generate realizations of synthetic connectomes from the ensemble of anatomically possible ones. Percentages of realizations in which each synapse is predicted correctly is a quality measure for synaptic specificity assumptions.

The first step defines all wiring diagrams that could in principle originate from the dense reconstruction (**Fig. 1b**). We generate such an *ensemble of connectomes*^5^ by modelling *possible synapses* as all combinations by which the pre- and postsynaptic sites in the data could connect to one another. If any presynaptic site could connect to any postsynaptic site, the number of connectomes in the ensemble is the *Factorial* of the number of synapses in the dataset (*n*_*syns*_). However, many of the combinatorially possible connectomes include connections between pre- and postsynaptic sites that are far apart. Users can exclude these anatomically implausible connectomes by setting a distance threshold beyond which connections are not possible. Here, we subdivide dense reconstructions into cubic subvolumes of 5, 10 or 25 µm edge lengths, and consider synapses only as possible if pre- and postsynaptic sites are within the same subvolume. Thus, we drastically reduce ensembles of combinatorially possible connectomes to anatomically plausible subsets – i.e., on the order of *Factorial*(*n*_*syns per subvolume*_). The first step enables statistical analysis of connectomes while accounting for the underlying dense anatomy.

The second step tests the null hypothesis that *“synaptic specificity is not the origin of the empirically observed connectivity patterns*”. For this purpose, we set the α_i_ parameters of all synaptic specificity assumptions to zero. Connectivity patterns derived from the resulting ensemble of connectomes could hence not originate from wiring that is conditional on neuronal properties. Instead, all combinations of connections between pre- and postsynaptic sites within a subvolume occur equally likely. Thus, the only source of wiring specificity is the morphology of the neurons, as it determines which pre- and postsynaptic sites are located in which subvolumes. In essence, the null hypothesis reveals the impact of neuron morphology on wiring specificity^5^. If the predicted connectivity patterns are consistent with those observed empirically, we cannot reject the null hypothesis. Thus, synaptic specificity is not required, and the impact of neuron morphology is sufficient, to account for these observations. In turn, if predicted and observed connectivity patterns deviate, we can reject the null hypothesis, and synaptic specificity is required to account for these observations. The second step hence distinguishes between connectivity patterns in dense reconstructions that could originate from morphological properties of the neurons and those that require additionally assumptions about synaptic specificity.

The third step tests the impact of synaptic specificity assumptions on wiring specificity (**Fig. 1c**). For this purpose, we consider α_i_ parameters as free model parameters and use a simulation-based Bayesian inference (SBI) algorithm that we developed^29^ to identify all α_i_ ∈ [−1, +1] values that account best for the observed connectivity patterns. As for the null hypothesis, two scenarios arise. If predicted and observed connectivity patterns match, the synaptic specificity assumptions are sufficient to account for these patterns. Thus, the corresponding inferred α_i_ values provide quantitative predictions for each observed synapse – and each possible but unobserved synapse – how negative or positive wiring preferences conditional on neuronal properties could explain these patterns. In turn, if predicted and observed connectivity patterns do not match, the synaptic specificity assumptions are not sufficient to account for these patterns. Thus, additional or different assumptions are required. Therefore, we enable users to test different assumptions separately and combinations of assumptions jointly. The third step hence reveals which synaptic specificity assumptions are sufficient to account for which connectivity patterns in dense reconstructions.

The fourth step evaluates the impact of synaptic specificity assumptions on each individual synapse in the data (**Fig. 1d**). At this point, we emphasize an important feature of our statistical approach. Synaptic specificity assumptions do not affect which connectomes could occur in principle, as the ensemble of connectomes is defined solely by the anatomy of the dense reconstruction. Instead, synaptic specificity assumptions affect how likely each anatomically possible connectome occurs. For example, in the null model – α_i_ = 0; *for all* i – all pre- and postsynaptic sites in a subvolume are equally likely to connect to one another. Therefore, the neuronal structures that contribute most pre- and postsynaptic sites to a subvolume are most likely to connect to one another, and hence, the corresponding connectome occurs most likely. In turn, if synaptic specificity parameters deviate from zero, the neuronal structures that contribute the most pre- and postsynaptic sites may no longer connect most likely to one another, and hence, a connectome other than the one of the null model may occur most likely. Thus, our approach enables us to quantify via a *loss function* how the likelihood for each possible synapse to occur changes across different synaptic specificity assumptions. The fourth step hence provides a measure of which assumptions could account best for each observed synapse in dense reconstructions.

The fifth step assess to what extent the inferred α_i_ values – and hence the corresponding synaptic specificity assumptions – account for the entire empirically observed connectome. For this purpose, we generate discrete realizations of synthetic connectomes from the ensemble of anatomically possible ones (**Fig. 1e**). The inferred α_i_ values would be ideal, if any realization that is generated with these values matched the empirically observed connectome. Therefore, we use the percentage of realizations in which each synapse is predicted correctly as a quality measure for synaptic specificity assumptions.

Ultimately, our workflow concludes by providing values for biologically interpretable assumptions that directly generate the empirically observed connectome from the underlying dense reconstruction.

### Dissecting origins of wiring specificity in mouse primary visual cortex

We demonstrate each step of our workflow using a recently reported dataset^27^ from the mouse primary visual cortex (V1). This dataset provides two inputs to our framework. First, a densely reconstructed volume of 250μm x 140μm x 90μm from the superficial layers (L2/3), which contains neuron somata, dendrites and axons, and 3.2 million synapses between them (**Fig. 2a-c**). Second, a connectome, which specifies the locations of all synapses between the neurons whose somata are located inside the densely reconstructed volume (**Fig. 2d-f**). As a third input, we selected four types of connectivity patterns whose deviations from a randomly connected network were considered as evidence for synaptic specificity ^1, 4, 9–14, 27^: whether a synapse occurs on an EXC or INH neuron (**Fig. 3a**), whether a synapse occurs on a soma, dendrite or AIS (**Fig. 3b**), how many synapses connect two neurons (**Fig. 3c**), and the motif that three neurons form (**Fig. 3d**). As a fourth input, we defined three synaptic specificity assumptions (**Fig. 4a**), which model that neurons may wire conditional on their cell types (here: EXC vs. INH), cellular properties (here: pairwise correlations), or subcellular properties (here: postsynaptic target sites). Based on these four inputs, our demonstration reveals which assumptions could account best for which observed synapses in this cortical dataset (**Fig. 5**), and how reliably each observed (and unobserved) synapse can be predicted from the dense reconstruction (**Fig. 6**).

**Figure 2:**
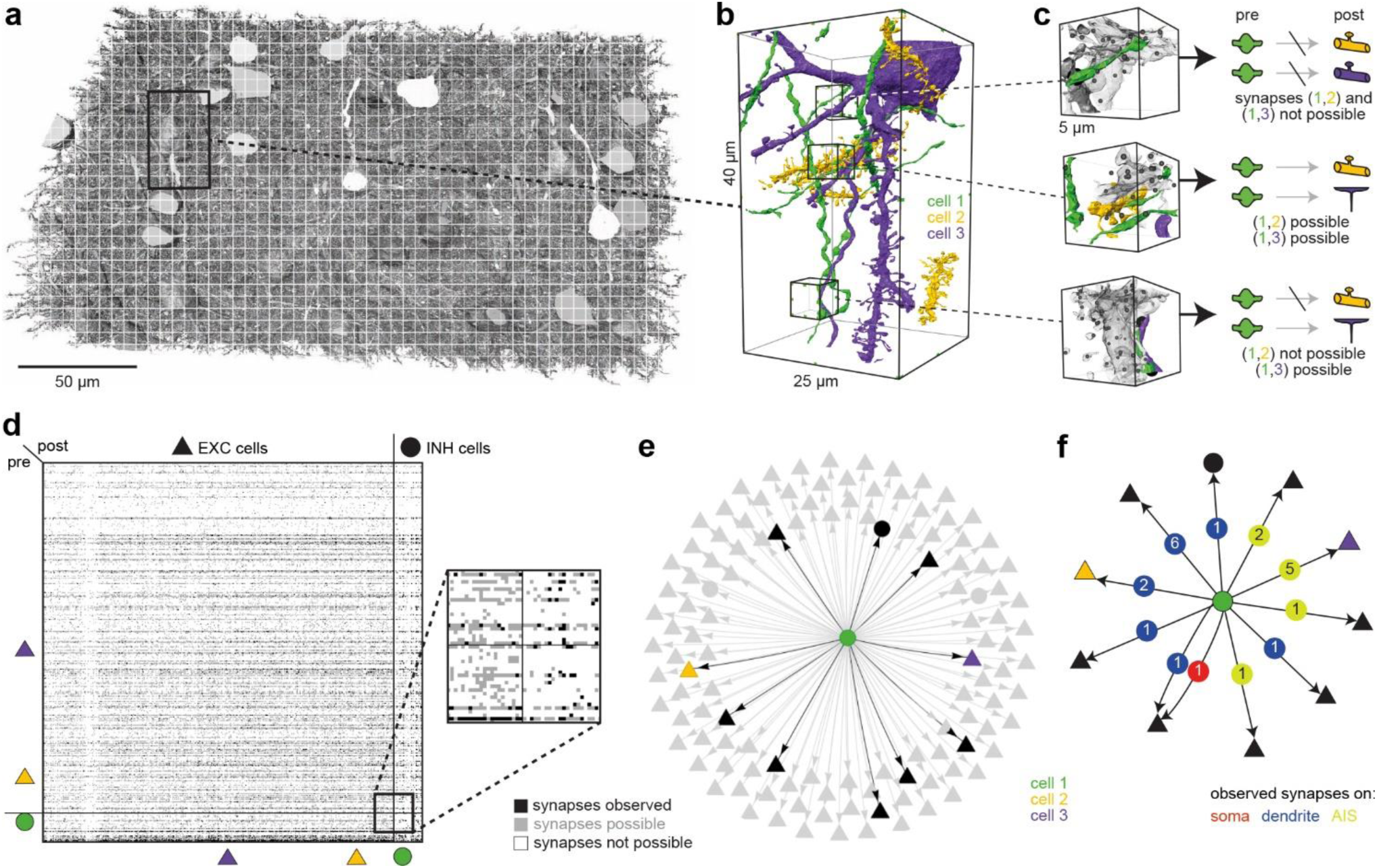
Step 1 – Ensembles of connectomes for a densely reconstructed volume of mouse V1. **(a)** We demonstrate our workflow on a dense dataset from mouse V1^27^. We partitioned the densely reconstructed volume uniformly into 5μm cubes (indicated by the white grid). **(b)** Zoom-in to panel a shows neuronal structures of three neurons whose somata are located in the dataset: axon segments of an inhibitory (INH) neuron (green), dendrite segments of an excitatory (EXC) neuron (yellow), and the soma, dendrite and the axon initial segments (AIS) of another EXC neuron (purple). **(c)** Zoom-ins to panel b show three examples of 5μm cubes. We consider synapses as possible if the observed pre- and postsynaptic structures are located in the same cube. Top: the INH neuron cannot connect to either of the two EXC neurons, because they have no postsynaptic structures in this cube. Center: presynaptic structures of the INH neuron could connect to postsynaptic structures on the dendrites (yellow) of the first and to postsynaptic structures on the AIS (purple) of the second EXC neuron. Bottom: presynaptic structures of the INH neuron could connect to postsynaptic structures on the AIS (purple) of the second EXC neuron, but not to the first EXC neuron. **Please note:** even though they are not shown in these examples, we consider all pre- and postsynaptic structures in all subvolumes of the dataset. **(d-f)** Visualizations of the ensemble of *anatomically possible* connectomes for 5μm grids. The anatomically possible connectomes (grey) include the empirically observed one (black). **(d)** Connectivity matrix for the 451 neurons whose somata are located within the reconstructed volume, grouped by EXC (triangle) and INH (circle) neurons. The example neurons from panels b-c are denoted by their respective colors. The zoom-in highlights that the number of neurons that could connect to one another (grey) exceeds the observed number (black), but that the number of neurons that could not connect to one another (white) is by far the largest group. **(e)** Node-link diagram for the example INH neuron (green) shows all the neurons it could connect to (grey) vs. those it was found to connect to (black). **(f)** The edges of the node-link diagram in panel e represent both the numbers and subcellular locations (on the soma, dendrite or AIS) of all possible and observed synapses.

**Figure 3:**
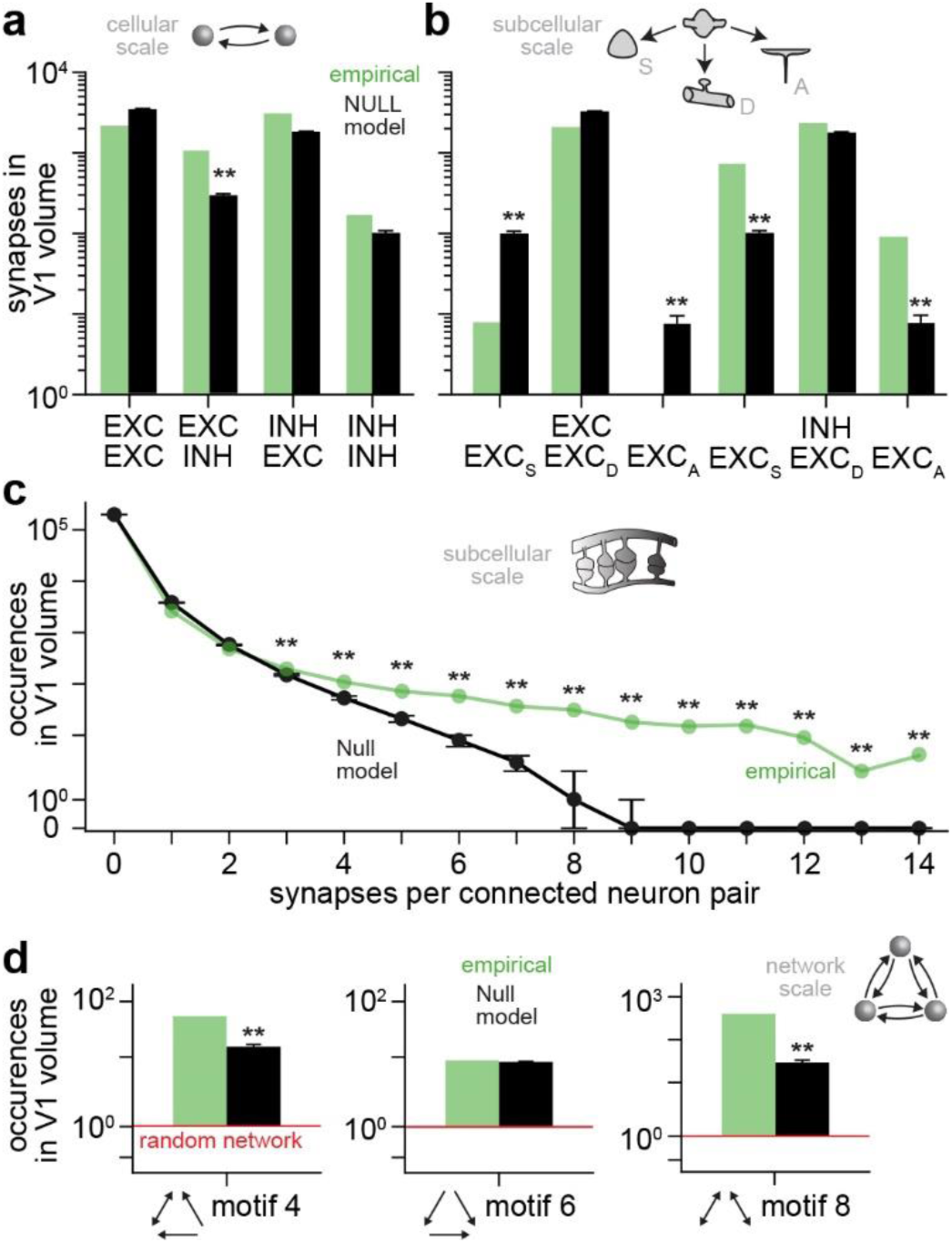
Step 2 – Testing the impact of neuron morphology on wiring specificity in mouse V1. **(a-d)** We tested the *null hypothesis* that synaptic specificity is not the origin for empirically observed wiring specificity on four types of connectivity patterns – i.e., we predicted these connectivity patterns under the assumption that wiring is not conditional on neuronal properties (all pre- and postsynaptic sites within a 5μm cube are equally like to connect to one another). This null hypothesis reveals the impact of neuron morphology on wiring specificity. We show the empirical observations in green, and the model predictions in black. The asterisks denote connectivity patterns that violate the null hypothesis – i.e., these observations reflect synaptic specificity. In turn, for patterns without asterisks, we could not reject the null hypothesis – i.e., these observations could reflect the impact of neuron morphology. **(a)** Observed occurrences of EXC synapses on INH neurons cannot be explained by the impact of neuron morphology. **(b)** Observed occurrences of EXC and INH synapses on EXC somata and AISs cannot be explained by the impact of neuron morphology. **(c)** Observed occurrences of more than three synapses per neuron pair cannot be explained by the impact of neuron morphology. **(d)** Observed occurrences of motif 4 and 8 cannot be explained by the impact of neuron morphology. See **Fig. S1** for more comparisons with additional connectivity patterns and for grid sizes of 10 and 25μm.

**Figure 4:**
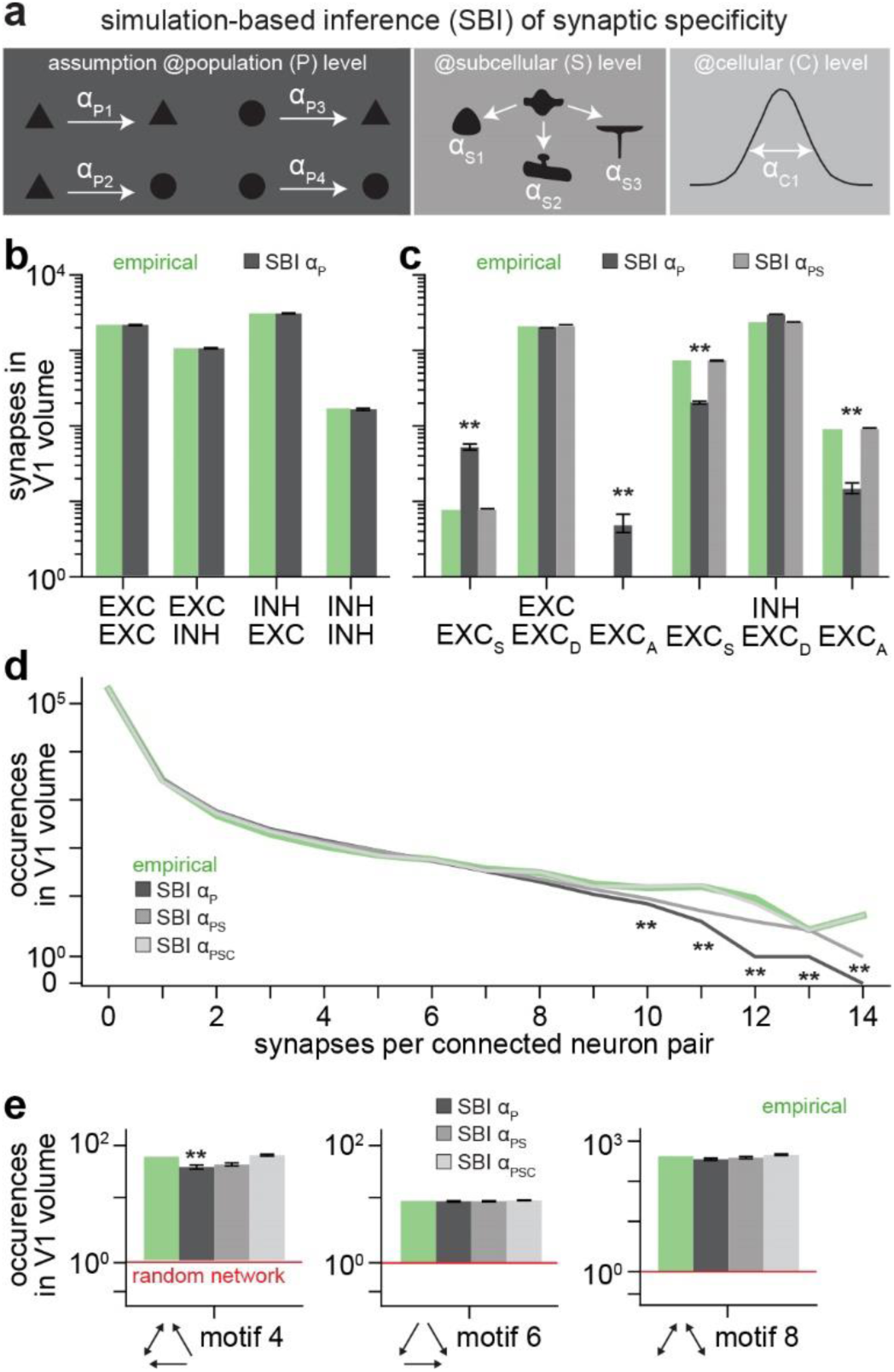
Step 3 – Testing the impact of synaptic specificity on wiring specificity in mouse V1. **(a)** We tested combinations of three synaptic specificity assumptions on the four types of connectivity patterns shown in Fig. 3. Left: wiring is conditional on whether pre- and postsynaptic sites belong to EXC or INH neurons – i.e., we define synaptic specificity parameters at population level α_*P*_. Center: wiring is conditional on the postsynaptic target site – i.e., we define synaptic specificity parameters at subcellular levels α_*S*_. Right: wiring is conditional on pairwise correlations – i.e., we define synaptic specificity parameters at cellular levels α_*C*_. **(b-e)** We used SBI to identify all α_i_ ∈ [−1, +1] values that accounted best for the observed connectivity patterns **(Fig. S5)**. We show the model predictions in grey shadings of the respective assumptions from panel a. The asterisks denote connectivity patterns that were not explained by the tested assumptions – i.e., additional or different assumptions would be required. **(b)** Observed occurrences of EXC and INH synapses could be explained by the impact of morphology in combination with wiring conditional on cell type (i.e., α_*P*_). **(c)** Explaining observed occurrences of EXC and INH synapses on EXC somata and AISs additionally requires wiring conditional on the postsynaptic target site (i.e., α_*PS*_). **(d)** Explaining observed occurrences of more than nine synapses per neuron pair additionally requires wiring conditional on pairwise correlations (i.e., α_*PSC*_). **(e)** Observed occurrences of motifs are generally well explained by the impact of neuron morphology in combination with wiring conditional on cell type (i.e., α_*P*_). See **Fig. S2** for more comparisons with additional connectivity patterns.

**Figure 5:**
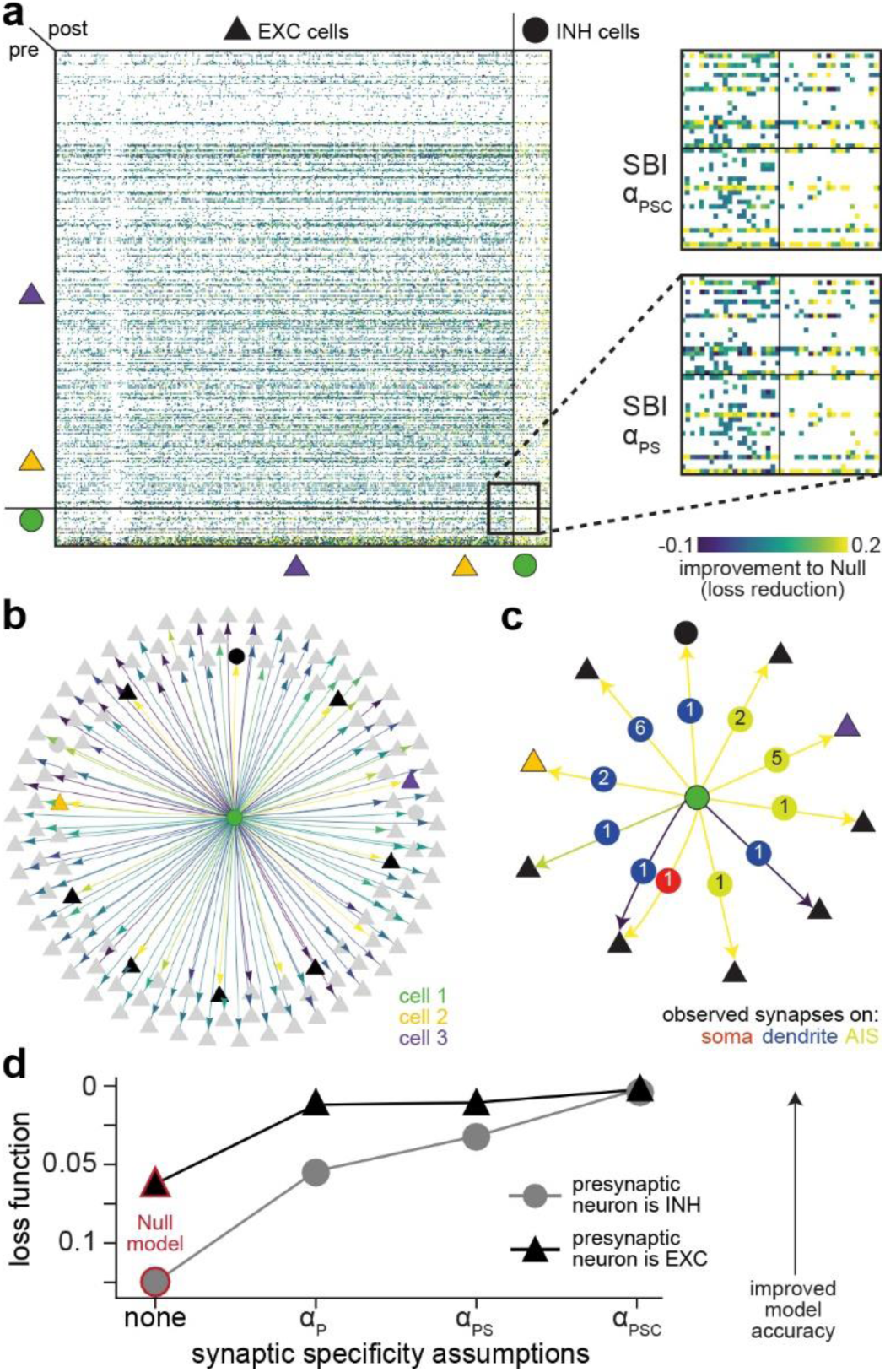
Step 4 – Testing the impact of synaptic specificity on each synapse in the V1 dataset. **(a-c)** We quantified how likely each possible synapse would occur under the null hypothesis, and how these likelihoods change for the different synaptic specificity assumptions in Fig. 4. We visualize the resultant loss functions for the ensemble of connectomes in Fig. 2, where now colors denote whether the tested synaptic specificity assumptions improved (yellow) or not improved (blue) the predictability of an observed synapse – and possible but unobserved synapse – compared to the Null model. **(a)** Left: combining the assumptions that wiring is conditional on EXC and INH cell types and on postsynaptic target site (i.e., α_*PS*_) increased the predictability for most synapses onto INH neurons, but increased the predictability much less for most synapses onto EXC neurons. Right: the zoom-ins show that adding the assumption that wiring is conditional on pairwise correlations (i.e., α_*PSC*_) did further improve the predictability of some synapses, primarily for those onto EXC neurons. **(b)** Visualization of the loss function for the example INH neuron from Fig. 2, illustrates that the same assumption (here α_*PS*_) can affect synapses from the same presynaptic neuron differently for different postsynaptic neurons. **(c)** This is even the case for synapses on the same postsynaptic neuron depending on whether synapses were located on the soma, dendrites or AIS. **(d)** Improvement of the predictability of synapses versus the complexity of synaptic specificity assumptions. Our assumptions at cell type level improved the predictions for both EXC and INH synapses, whereas our assumptions at cellular and subcellular levels improved primarily the predictions for INH synapses.

**Figure 6:**
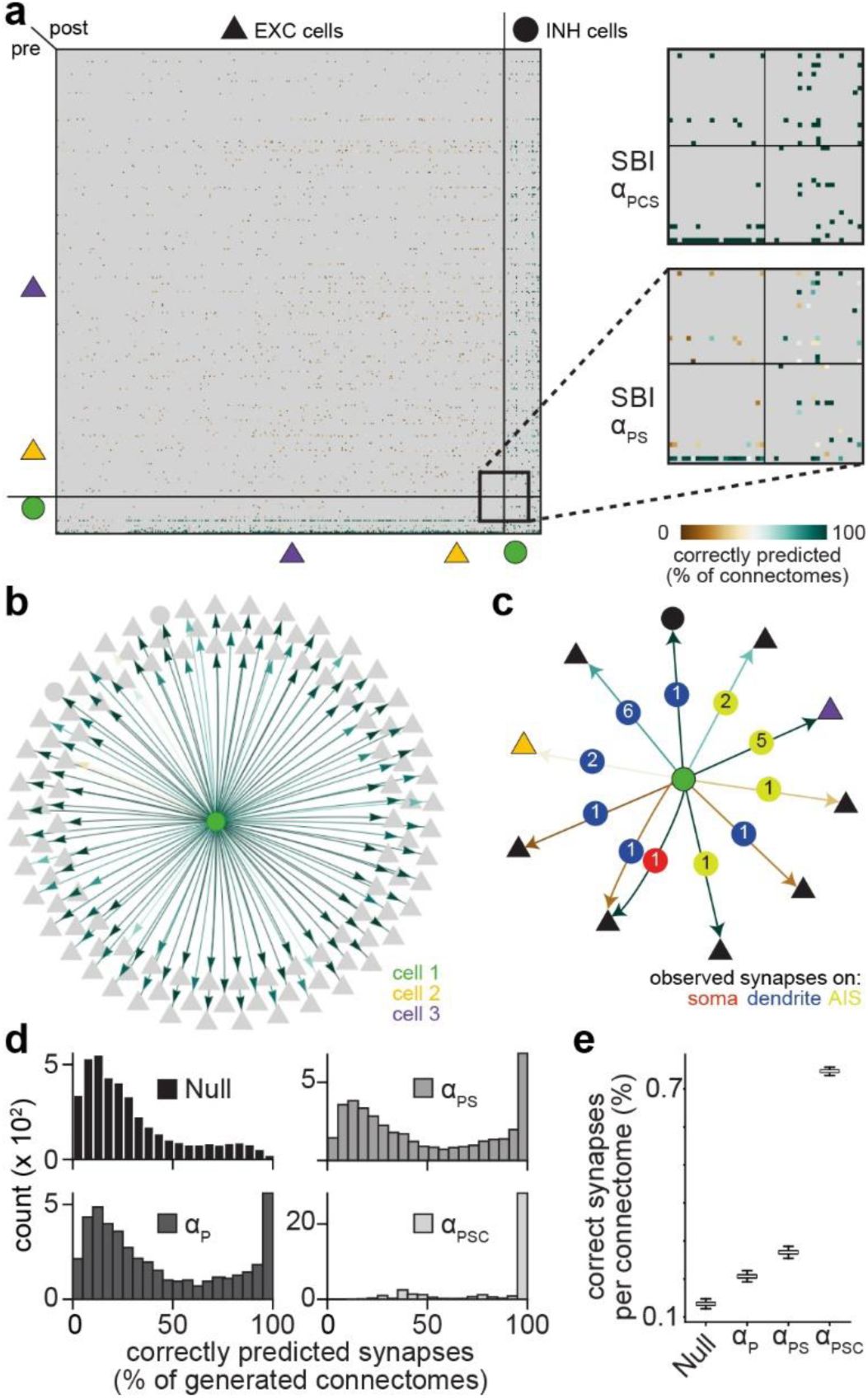
Step 5 – Testing the impact of synaptic specificity on the connectome from mouse V1. **(a-d)** We compared the empirically observed connectome with realizations that we generated from the ensemble of anatomically possible ones. We quantified how often each observed synapse occurs across 1,000 realizations that we generated with the same parameters. We visualize the resultant percentages for the ensemble of connectomes as in Fig. 5, but now the colors denote the fraction of realizations in which a synapse was correctly predicted. **(a)** Left: the assumptions that wiring is conditional on cell type and on postsynaptic target site (i.e., α_*PS*_) predicted reliably synapses from and onto INH neurons (turquoise), but generally failed to predict EXC-to-EXC synapses (brown). Right: the zoom-ins show that by adding wiring conditional on pairwise correlations (α_*PSC*_), virtually all synapses are predicted reliably, including EXC-to-EXC synapses. **(b)** Visualization of the ability to predict the synapses from the INH neuron from Fig. 2, illustrates that the same assumption (here α_*PS*_) can affect synapses from the same presynaptic neuron differently for different postsynaptic neurons. **(c)** This is also the case for synapses on the same postsynaptic neuron depending on whether synapses were located on the soma, dendrites or AIS. **(d)** Histograms of how often each of the empirically observed synapses was correctly predicted in 1,000 connectomes generated by the Null model (black) or by each combination of our synaptic specificity assumptions (grey shadings), respectively. The null hypothesis predicted synapses correctly in <1% of the connectomes. Our synaptic specificity assumptions increased the percentages of correct predictions to 13% (α_*P*_), 17% (α_*PS*_), and 71% (α_*PSC*_), respectively. **(e)** Correctly predicted synapses in each of the 1,000 connectomes that we generated respectively by the Null model or by each combination of our synaptic specificity assumptions. See **Fig. S3** for a grid size comparison.

#### Step 1

We generated the ensemble of connectomes for this dataset. We partitioned the cortical volume into 5μm cubes (**Fig. 2a-b**) and considered synapses as *possible* if the pre- and postsynaptic sites are located in the same cube, and as *not possible* otherwise (**Fig. 2c**). In each 5μm cube, we observed, on average, 100 synapses (32,438 cubes; median/25^th^/75^th^ percentile: 117/61/139). As the respective pre- and postsynaptic sites of these 100 synapses could, in principle, connect to one another, the number of *possible* connectomes with 100 synapses per 5μm cube is virtually infinite – *Factorial*(100) ≈ 10^158^. Even so, the number of connectomes that we excluded as anatomically *not possible* is much larger – i.e., *Factorial*(3.2×10^6^) ≫ *Factorial*(100). Thus, compared to generative network models that do not consider the dense reconstruction of neuron morphology^2^, our approach reduces the connectomes that could possibly arise for this cortical dataset to a drastically smaller, yet still enormous, subset. This subset of *anatomically plausible connectomes* included the empirically observed one (**Fig. 2d-f**).

#### Step 2

We tested the *null hypothesis* that synaptic specificity is not the origin of connectivity patterns in this cortical dataset. For this purpose, we predicted connectivity patterns under the assumption that wiring is not conditional on neuronal properties – i.e., all pre- and postsynaptic sites within a 5μm cube are equally likely to connect to one another. For several patterns, we found that the differences between model predictions and empirical observations were not sufficient to reject the null hypothesis (**Fig. 3a-d)**. The occurrences of INH synapses on EXC and INH neurons, of EXC and INH synapses on EXC dendrites, of up to three synapses per neuron pair, and of feedforward-feedback loops between three neurons (motif 6), were all predicted well without any synaptic specificity assumptions. As we showed previously^5^, the impact of neuron morphology could hence account for these connectivity patterns. In contrast, the model predictions for the remaining connectivity patterns violated the null hypothesis (**Fig. 3a-d**). The model predicted occurrences of EXC synapses on INH neurons, of EXC and INH synapses on somata and AISs, of more than three synapses per neuron pair, and of recurrent feedback loops (motif 4) differed significantly from the respective empirical observations. Thus, we demonstrate that our approach differentiates between connectivity patterns that could originate from morphological properties of the neurons and those that require additionally synaptic specificity assumptions.

#### Step 3

We tested whether our synaptic specificity assumptions could be an origin for the connectivity patterns that violated the null hypothesis. First, we assumed that wiring is conditional on whether pre- and postsynaptic sites belong to EXC or INH neurons (**Fig. 4a**). For this purpose, we defined synaptic specificity parameters at the population level α_*P*1−4_ (i.e., EE, EI, IE, II) and used SBI to identify the α_*P*1−4_ ∈ [−1, +1] values that accounted best for the observed connectivity patterns. Compared to the null hypothesis, this assumption improved the model predictions for all patterns substantially (**Fig. 4b-e**). In fact, for occurrences of EXC and INH synapses on EXC and INH neurons, and of the motifs in general, the differences between model predictions and empirical observations disappeared virtually completely. In contrast, predicted and observed occurrences of EXC and INH synapses on somata and AISs, and of more than nine synapses per neuron pair, still deviated significantly. However, it is worth noting that the occurrences of up to nine synapses per neuron pair were now well predicted. In essence, the assumption that neurons wire conditional on cell types (i.e., EXC and INH) is sufficient to account for nearly all cellular and network scale connectivity patterns that we tested in this cortical dataset.

While accounting for nearly all cellular and network scale connectivity patterns, our assumption that wiring is conditional on cell type did generally not account for subcellular scale connectivity patterns. Therefore, we tested the additional assumption that wiring is conditional on the postsynaptic target site (**Fig. 4a**). For this purpose, we defined three synaptic specificity parameters at subcellular levels α_*S*1−3_ (i.e., onto soma, dendrite, AIS). Thus, in combination with our first assumption, we now used SBI to identify which values of twelve parameters α_*PS*1−12_ ∈ [−1, +1] – i.e., three α_*PS*_ parameters for EE, EI, IE, and II – accounted best for the observed connectivity. The combination of these two assumptions improved the model predictions further (**Fig. 4b-e**). In fact, differences between model predictions and empirical observations disappeared completely, except for occurrences of more than nine synapses per neuron pair. This small, but significant difference disappeared, however, when we included our third assumption that wiring is conditional on pairwise correlations (**Fig. 4a**). For this purpose, we defined synaptic specificity parameters at cellular levels α_*C*_. In combination with our first two assumptions, SBI now identified values that accounted for all empirically observed connectivity patterns (**Fig. 4b-e)**. Thus, by testing synaptic specificity assumptions from subcellular to population levels separately and jointly, we demonstrate that our approach can reveal which of these assumptions could be sufficient to account for which connectivity patterns in this cortical dataset.

#### Step 4

We evaluated the impact of our assumptions on each individual synapse in this cortical dataset. We quantified how likely each anatomically possible synapse would occur under the null hypothesis, and how these likelihoods change for different synaptic specificity assumptions (**Fig. 5a**). The resulting loss function revealed which observed synapses become more likely if wiring would be conditional on different neuronal properties, and which of the possible, but unobserved synapses become less likely. We found that each assumption can make some synapses more likely, others less likely, or not change their likelihood (**Fig. 5a-c**). In fact, the same assumption can affect synapses from the same presynaptic neuron differently for different postsynaptic neurons (**Fig. 5b**). This was even the case for synapses on the same postsynaptic neuron depending on whether synapses were located on the soma, dendrites or AIS (**Fig. 5c**). Moreover, we found that synaptic specificity assumptions impacted the predictability of INH synapses more than those of EXC synapses (**Fig. 5d**). Our assumptions at cell type level improved the model predictions for both EXC and INH synapses, whereas our assumptions at cellular and subcellular levels improved primarily the predictions for INH synapses. In essence, EXC connectivity patterns in this cortical dataset can be predicted with less (complex) assumptions than those with INH synapses. Overall, we demonstrate that our approach reveals which synaptic specificity assumptions could account best for each observed synapse in a dense reconstruction.

#### Step 5

We tested to what extent our synaptic specificity assumptions could account for the entire connectome from this cortical dataset. For each assumption, we compared the observed connectome with 1,000 realizations generated from the anatomically possible ensemble (**Fig. 6a**). We hence quantified how often each observed synapse occurred across connectomes generated with the same parameters. In line with our loss function analysis, we found that the same assumption can affect synapses differently, both from the same presynaptic neuron (**Fig. 6b**) and between the same pre- and postsynaptic neurons (**Fig. 6c**). In fact, the same assumption may predict the synapses that a neuron forms on the soma in all generated connectomes, whereas the synapses it forms on the dendrites of this neuron are never predicted correctly. Also consistent with our loss function analysis, the ability to predict synapses increased with the number of assumptions (**Fig. 6d**). Under the null hypothesis, each synapse was predicted correctly in <1% of the 1,000 connectomes. Generating connectomes without synaptic specificity assumptions will hence lead virtually never to the observed connectome. In essence, neuron morphology is insufficient to predict any of the synapses in this dataset. However, by incorporating synaptic specificity at cell type, and additionally, at subcellular and cellular levels, each synapse was predicted correctly in 13%, 17% and 71% of the connectomes, respectively. In fact, any of the 1,000 connectomes generated with just three synaptic specificity assumptions predicted 75% of the connectome as observed empirically at nanoscale resolution (**Fig. 6e**). Overall, this result indicates that our method can predict values for biologically interpretable synaptic specificity parameters that generate the empirically observed connectome directly from the underlying dense reconstruction.

We provide the predicted wiring preferences of each observed synapse as the set of α ∈ [−1, +1] values that SBI inferred for the different synaptic specificity parameters. In essence, these sets provide the predictions for the respective negative and positive wiring preferences that account for each of the observed synapses in the dense reconstruction. This unique information at synaptic resolution opens new avenues for research, which we illustrate by two examples: neuron classification based on synaptic specificity and by comparing synaptic specificity predictions across datasets.

### Perspective 1: Neuron classification based on synaptic specificity

The mouse V1 dataset comprises 417 EXC (i.e., ‘pyramidal’) and 34 INH neurons, whose somata are contained within the reconstructed volume. Based on morphological properties^27^, 20 INH neurons are annotated as ‘bipolar’, ‘basket’, ‘chandelier’, ‘martinotti’ or ‘neurogliaform’ cells. To complement this morphological classification, we visualized the neurons in the principal component (PC) space that is based on their respective set of synaptic specificity values (**Fig. 7a**). The EXC neurons clustered within a narrow band of PC1, whereas the INH neurons spread across the range of both PC1 and PC2. INH neurons of the same class were, however, closer to one another in the synaptic specificity space than those of different classes (**Fig. 7b**). Indeed, clustering based on synaptic specificity identified three clusters that separated remarkably well between chandelier, basket and bipolar cells (**Fig. 7c**). These clusters resulted primarily from differences at subcellular level (**Fig. 7d-e**). SBI predicts that chandelier cells have strong positive preferences to wire on AISs of EXC neurons, but negative preferences to wire on EXC dendrites. In contrast, bipolar and basket cells have positive preferences to wire on both EXC somata and AISs, but basket cells have, in addition, negative preferences to wire on EXC dendrites. Notably, SBI predicted that both EXC and INH neurons have no preferences for targeting specific domains of INH neurons (**Fig. 7e**). These results indicate substantial differences for how EXC and INH neurons wire conditional on their neuronal properties, that INH cell types differ in how they wire conditional on target domains, but that these differences are limited to INH-to-EXC synapses.

**Figure 7:**
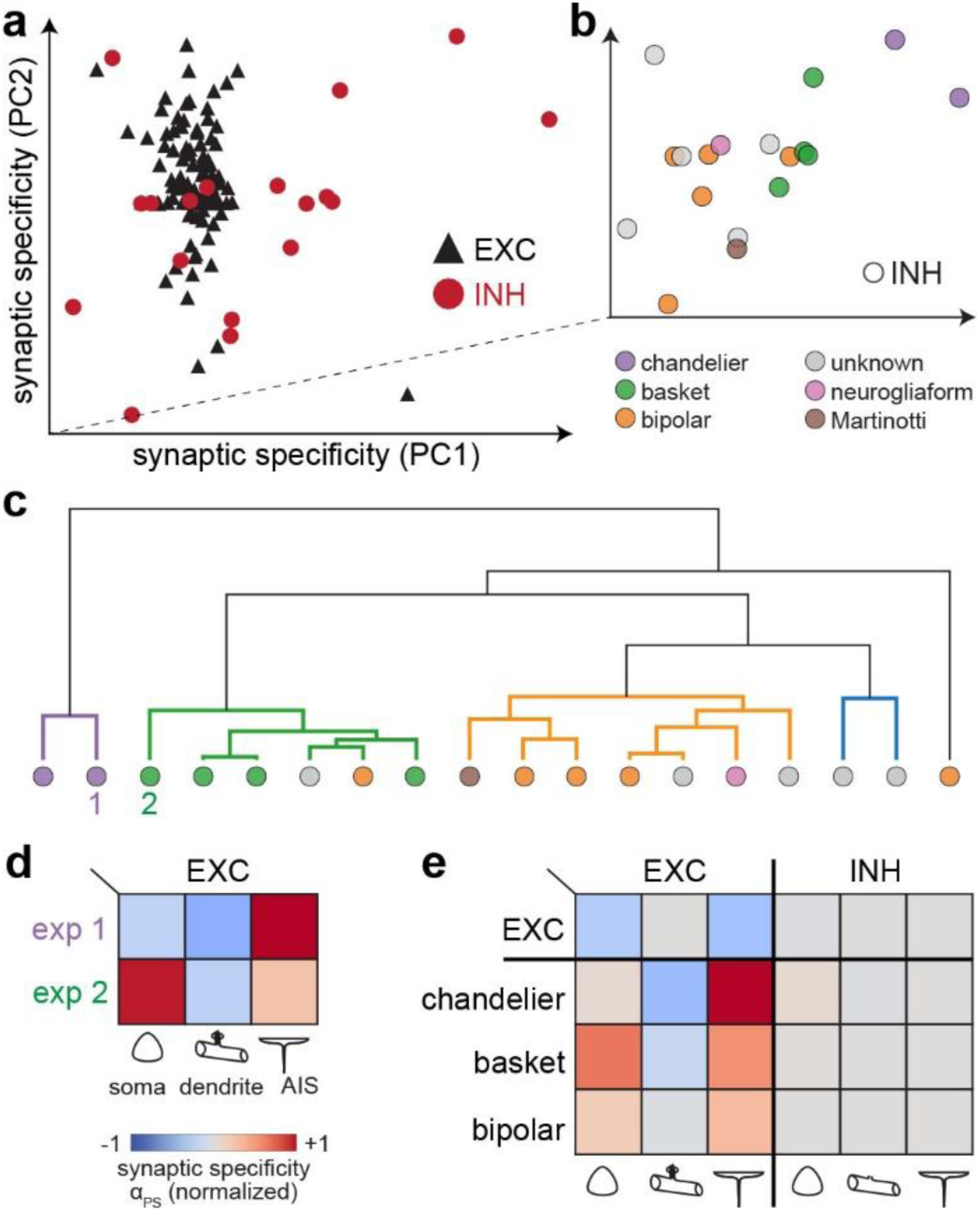
Neuron classification based on synaptic specificity. **(a)** The mouse V1 dataset comprises 417 EXC and 34 INH neurons, of which 20 are further subclassified as ‘bipolar’, ‘basket’, ‘chandelier’, ‘martinotti’ or ‘neurogliaform’ cells. We visualized 127 neurons (109 EXC and 18 INH) with at least 3 outgoing synapses in the principal component (PC) space that is based on their respective set of inferred synaptic specificity values. **(b)** INH neurons of the same subclass were closer to one another in the synaptic specificity space than those of different classes. **(c)** Hierarchical clustering based on synaptic specificity identified three clusters that separated remarkably well between chandelier, basket and bipolar cells. **(d)** SBI predicts that chandelier, basket and bipolar cells have different preferences to wire on somata, dendrites and AISs of EXC neurons. **(e)** SBI predicts that both EXC and INH neurons have no wiring preferences for targeting specific domains of INH neurons.

### Perspective 2: Comparing synaptic specificity predictions across datasets

We compared our results for mouse V1 with those for a recently reported dataset from human temporal cortex^14^. The human dataset comprised a much larger densely reconstructed volume of ∼1mm^3^, which contained 133.7 million synapses (**Fig. 8a**). We analyzed the human dataset analogously to the mouse dataset – i.e., we selected the same four types of connectivity patterns for analysis and tested the same three synaptic specificity assumptions (**Fig. 8b-e**). The results from each step of our workflow were remarkably similar between both datasets. Morphological properties of the neurons in both datasets were sufficient to predicted the occurrence of INH synapses on EXC and INH neurons (**Fig. 8b**), of EXC and INH synapses on EXC dendrites (**Fig. 8c**), of multiple synapses between neuron pairs (**Fig. 8d**), and of feedforward-feedback loops (motif 6) (**Fig. 8e**). Also consistent between the two datasets, the assumption that neurons wire conditional on cell types (i.e., EXC and INH) could account for nearly all cellular and network scale connectivity patterns that we tested, but could generally not account for subcellular scale connectivity patterns. In both datasets, the assumption that wiring is conditional on postsynaptic target sites was necessary to predict the observed occurrences of synapses on the somata, dendrites and AISs, while the assumption that wiring is conditional on pairwise correlations was necessary to predict the occurrences of multi-synaptic connections. These results indicate that the same synaptic specificity assumptions are sufficient to account for the same types of connectivity patterns in these very different cortical datasets.

**Figure 8:**
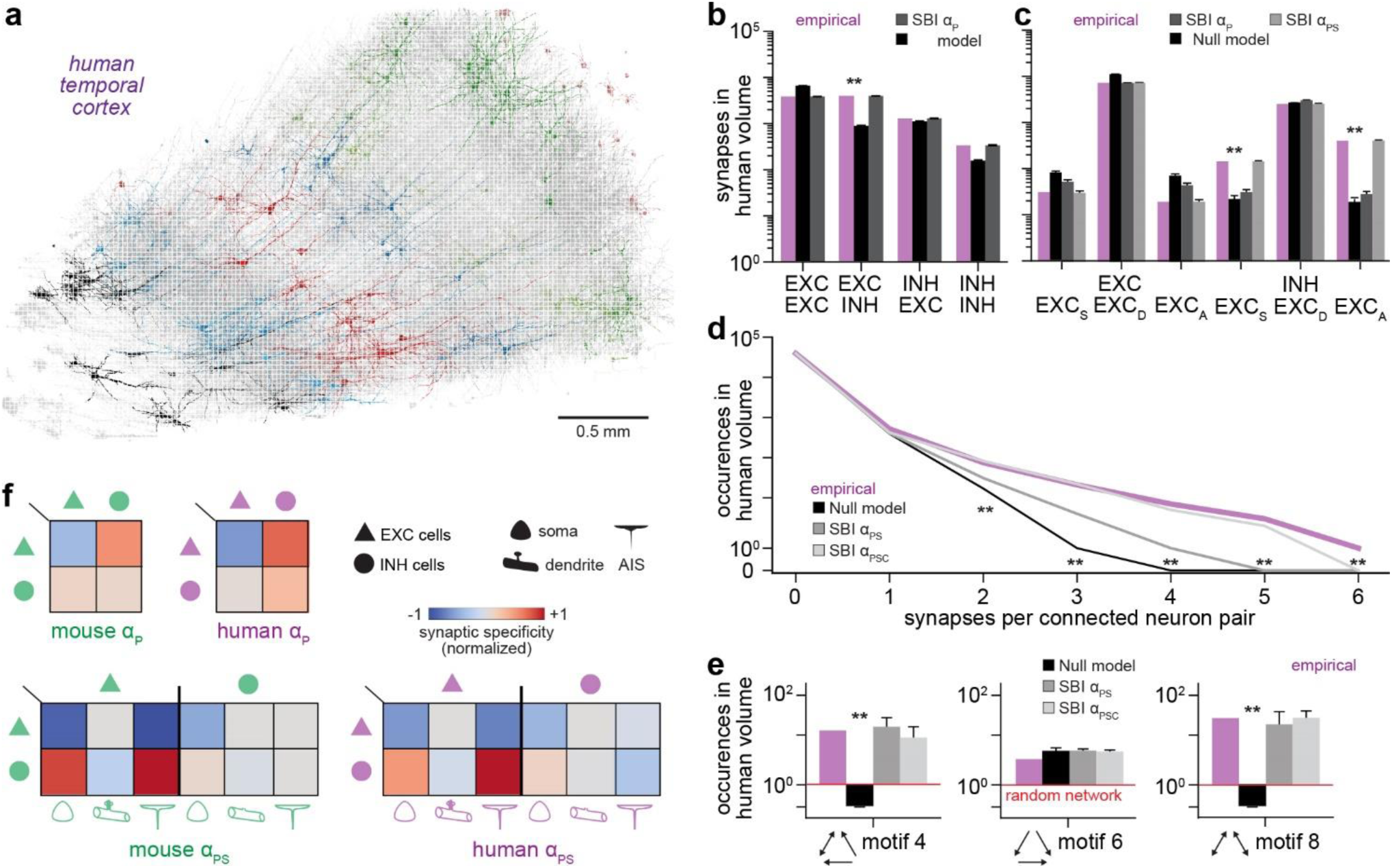
The origins of wiring specificity in mouse V1 versus human temporal cortex. **(a)** We repeated our analysis from Fig. 2**-6** for a densely reconstructed volume from human temporal cortex^14^. **(b-e)** We analyzed the human dataset analogously to the mouse dataset – i.e., we analyzed the same types of connectivity patterns and tested the same synaptic specificity assumptions. The results indicate that the same assumptions are sufficient to account for the same types of connectivity patterns also in this very different cortical datasets. **(f)** α ∈ [−1, +1] values that our workflow inferred for the mouse (green) and human (purple) datasets. Top panels: SBI predicted for both datasets that EXC neurons have negative preferences to wire on EXC neurons, but positive preferences to wire on INH neurons. Bottom panels: SBI predicted for both datasets that EXC neurons have strong negative preferences to wire on EXC somata and AISs, whereas INH neurons have strong positive preferences for these EXC target domains. In contrast, neither EXC nor INH neurons in both datasets have wiring preferences for EXC and INH dendrites, or for targeting specific domains of INH neurons in general. See **Fig. S4** for additional connectivity patterns in the human temporal cortex.

Most noticeably, also the α ∈ [−1, +1] values that our workflow inferred for the synaptic specificity parameters were qualitatively, and even quantitatively, similar between the two datasets (**Fig. 8f**). SBI predicted for both datasets that EXC neurons have negative preferences to wire on EXC neurons, but positive preferences to wire on INH neurons. Notably, for both datasets, wiring preferences from INH neurons onto both EXC and INH neurons were much less pronounced. These similarities generalized to the subcellular level. In both datasets, EXC neurons have strong negative preferences to wire on EXC somata and AISs, whereas INH neurons have strong positive preferences for these EXC target domains. In contrast, neither EXC nor INH neurons in both datasets have wiring preferences for EXC and INH dendrites, or for targeting specific domains of INH neurons in general. These results indicate that the same synaptic specificity assumptions, with nearly identical parameter values, may generate empirically observed connectomes across brain areas and species.

## Discussion

We introduce a method that reveals for each synapse in densely reconstructed volumes, which synaptic specificity assumptions are necessary, sufficient, and best-suited to account for it. We demonstrate the applicability of our method on two recently reported datasets from mouse primary visual cortex^27^ and human temporal cortex ^14^. We consider these demonstrations by no means as a final assessment on the origins of wiring specificity in these cortical areas. The dataset from mouse cortex comprises only a relatively small volume, the dataset from human cortex still lacks final proof-reading, and we tested only three assumptions about synaptic specificity. However, despite these limitations, the demonstrations show that our method provides already at this stage some remarkable insights. First, while we confirm our previous finding^5^ that the impact of neuron morphology could account for non-random wiring at network scales (e.g. motif occurrences), we find that it does not account for any of the observed synapses in these datasets. Second, a small set of just three assumptions accounted for 75% of the connectome from mouse V1. Third, these three assumptions, with virtually the same values for negative or positive wiring preferences, accounted for connectivity in both the mouse and human datasets. We discuss implications of these striking results in separate sections below.

Before proceeding further, we will discuss some of the technical features of the method we employed. Our method seeks to identify a minimal set of assumptions that generate synthetic connectomes which closely resemble the properties of empirically observed connectomes. From a technical point of view, our method hence falls into the category of generative network models^25, 30^. Generative network models have been used successfully to model connectomes at both the microscale^2, 3^ and macroscale^25, 31–33^. At microscales, nodes and edges of the generated networks represent point neurons and synaptic connections between them. At macroscales, the nodes and edges represent brain areas and structural or functional connectivity measures between them that are derived from MRI data. For example, Haber et al. showed that by constraining the generation of microscale networks to match simple connectivity features observed in dense reconstructions – e.g. reciprocity of connectivity – the resultant synthetic connectomes replicate many other of the observed connectivity features – e.g. motif occurrences^2^. Beul et al. had similar results for macroscale network models by constraining their generation with simple features observed in MRI data – e.g. presence or absence of inter-areal connections^32^.

Our method goes beyond these state-of-the-art examples for generative modelling in five major ways. First, we generate connectomes at the nanoscale – i.e., we generate networks that replicate synapse locations on neuronal structures as observed at electron-microscopic resolution. Second, we generate connectomes directly from dense reconstructions. Thus, our generative network modelling accounts for the morphological properties of the neurons, and reveals how these properties impact connectivity^5^. Third, we generate connectomes based on parameters that reflect biologically interpretable synaptic specificity assumptions for how neurons wire conditional on their properties^17–19^. Thus, all connectivity features are model predictions, none are used as constraints for model generation (e.g. reciprocity or cell type connectivity^2^). Fourth, we use SBI^29^ to identify the distributions of all synaptic specificity values that can generate, equally well, the empirically observed connectome. Finally, for each synapse that is observed in dense reconstructions, we provide measures that quantify which assumptions are necessary, sufficient and best-suited to generate it. Taken together, our method extends beyond current generative network modelling approaches as it provides experimentally testable predictions of wiring preferences from subcellular to cell type levels that account for each synapse in dense reconstructions.

### Neuron morphology and synaptic specificity impact connectivity at different scales

How the morphological properties of the neurons impact their connectivity has been controversial for decades ^20^. This controversy results primarily from the common assumption that the impact of neuron morphology could be assessed by testing “Peters’ rule”^26^. According to this rule, axons form synapses randomly wherever they get close to a dendrite^34^. Peters’ rule may hence be simply stated as “proximity predicts connectivity”^5^. However, we had previously shown that proximity could predict connectivity in principle for only a very small minority (<1%) of the neuronal structures^5^. In essence, the number of axon-dendrite branch pairs in any subvolume (e.g. 5µm cube) of the cortex generally exceeds the number of synapses in this subvolume by one to two orders of magnitude. The vast majority of axon-dendrite branch pairs in any small cortical subvolume will hence always be unconnected, despite their proximity. Therefore, and in contrast to a widely held belief, observations that violate Peters’ rule^1, 8–14, 20, 35^ provide no evidence that these connectivity patterns result from synaptic specificity – i.e., for mechanisms that wired these neuronal structures conditional on their properties. In fact, virtually any observation that violates Peters’ rule may arise from our null hypothesis that *“synaptic specificity is not the origin of empirically observed connectivity patterns”*. Thus, testing Peters’ rule is not equivalent to testing the impact of neuron morphology on connectivity^5^. Instead, the impact of neuron morphology on connectivity can be revealed by testing our null hypothesis on dense reconstructions via our generative network modelling method – i.e., we test whether the morphological properties of the neurons account for the observed connectivity patterns (and each observed synapse), or whether additional assumptions about synaptic specificity are required for this.

We previously showed that the impact of neuron morphology leads unavoidably to wiring specificity from subcellular to network scales^5^. Network models that are generated from dense reconstructions under our null hypothesis – i.e., that all pre- and postsynaptic structures within a subvolume are equally likely to connect to one another – will hence generally display multi-synaptic connections and clusters of synapses, cell type- and location-specific connectivity, and non-random network motifs. In contrast, network models that are generated by the similar null hypothesis that all neurons in the entire volume are equally likely to connect to one another, and which hence neglects the morphological properties of the neurons, will generally not display any wiring specificity^2^. Therefore, connecting point neurons randomly yields random networks, whereas connecting real neurons randomly in accordance with their morphological properties yields ‘nonrandom’ networks – i.e., they have different connectivity patterns from subcellular to network scales^5^. In essence, limiting the ensemble of combinatorially possible connectomes to subsets of anatomically plausible ones is necessary, but also sufficient, to reveal the impact of neuron morphology on connectivity. Thus, generative network models at microscales will always require assumptions to account for the impact of neuron morphology on connectivity, whereas our approach reveals this impact in parameter-free network models. Taken together, we conclude that generative models may only be able to uncover the principles by which a neural network is organized if they take the morphological properties of its neurons into account.

The results from the demonstrations of our method on the dense datasets from mouse and human cortex support this conclusion. In line with our previous reports^5, 29, 36^, we found also for these datasets that the morphological properties of the neurons yield connectivity patterns at the network scale that are similar to those observed empirically. In fact, neuron morphology predicted some of the observed connectivity patterns perfectly, in particular non-random occurrences of network motifs. As a result, any network that is realized from anatomically plausible subsets of connectomes – without additional parameters and assumptions – will share most of the complex topological properties of the observed connectome. In essence, neuron morphology restricts the possible space of connectomes to those that share topological features of the observed connectomes. However, our demonstration also shows that none of the hence generated networks could capture the observed connectivity patterns at cellular and subcellular scales. In fact, without synaptic specificity assumptions, none of the generated networks could reliably predict the numbers and locations of synaptic connections between any neurons in these datasets. Thus, neuron morphology could account for topological properties of these cortical networks, but synaptic specificity was generally needed to predict connectivity at nanoscales. This result suggests that the origins of network topology – e.g. neuron morphology – are ‘decoupled’ from those that shape cell-to-cell connectivity – i.e., synaptic specificity. In any event, our results emphasize that generative network models need to consider both synaptic specificity and neuron morphology in order to capture the origins of wiring specificity in the cerebral cortex.

### Perspectives: Testing origins of wiring specificity across brain areas, species and time points

A striking result of our method demonstration is the finding that a just three simple assumptions about synaptic specificity were sufficient to predict the vast majority of synapses in the mouse dataset. Even more striking is our finding that the very same assumptions, with nearly identical parameter values for negative and positive wiring preferences, could account for the connectivity patterns in both the mouse and human datasets. Given the limitations discussed above, we will, however, not draw conclusions from these results. Instead, we consider them as an illustration for perspectives that our method offers.

Our method provides quantitative predictions for negative and positive preferences by which neurons establish synaptic connections with one another. Moreover, we demonstrate that our method enables to compare these predictions across neurons in a dataset, but also across datasets. For example, we predict that INH neurons in mouse visual cortex have similar wiring preferences if they are of the same morphologically determined cell type. Moreover, we predict that EXC neurons in both mouse visual and human temporal cortex have a lower preference to connect to EXC neurons than INH neurons, whereas INH neurons have a higher preference to connect to EXC neurons than INH neurons. Such quantitative predictions for well-defined cell types will enable experiments that investigate the molecular basis of these wiring preferences^15–18^. In fact, a recent study demonstrates the ability to link neurons from dense reconstructions to transcriptomics data^37^. Furthermore, these experiments could be combined with comparisons of the same brain area across individuals, for example to investigate how the impact of neuron morphology – and of synaptic specificity assumptions – may change during different stages of learning, development or disease^38^. Overall, our generative modelling method sets the stage to uncover the principles by which neural networks are organized from subcellular to network scales, and potentially to reveal the mechanisms by which they grow and develop.

## Materials and Methods

### Ensemble of anatomically possible connectomes (step 1)

The first input to our method is a dense reconstruction of neural tissue. Here, we analyzed a densely reconstructed volume from the mouse primary visual cortex (V1)^27^. We downloaded the synaptic connectivity information, the reconstructed meshes of the annotated neuron morphologies, and all corresponding metadata from https://doi.org/10.5281/zenodo.3710458. We also included annotations of synapses located on AISs as reported in a study from this dataset^13^. Synapses not located on AISs, were assigned to the postsynaptic compartment label ‘dendrite’. Synapses that were closer than 15μm to the center of the soma of the postsynaptic neuron were assigned to the postsynaptic compartment label ‘soma’. We partitioned the volume using 5 / 10 / 25 μm sized uniform grids resulting in 32441 / 4431 / 361 subvolumes. All synapses were assigned to one subvolume *k* based on their 3D locations. For this purpose, we used the mean position between the center locations of the pre- and postsynaptic structures of the synapse (e.g., bouton and dendritic spine) that were provided as annotations in the dense reconstruction dataset. Inside each subvolume, we processed the synapses *n*_*k*_ as follows: We split each synapse into a pre- and postsynaptic site and recorded for these sites the annotations of the corresponding neurons; the IDs of the pre- and postsynaptic neurons, their cell types and postsynaptic target labels (i.e., dendrite, soma, AIS). Pre- and postsynaptic sites that could not be associated with these annotations, e.g., due to incompletely traced axons or dendrites, were annotated as ‘unknown’ (i.e., −1). We refer to such neuronal structures that cannot be associated with a soma in the volume as ‘orphaned’ and to synapses from/to such neuronal structures as ‘unassigned’ synapses. We export the hence preprocessed synapses into csv-files (one for each grid size) with 7 columns: presynaptic neuron ID, postsynaptic neuron ID, presynaptic cell type, postsynaptic cell type, postsynaptic target site, subvolume ID, and the number of the respective synapses of this category. When the csv-file for a given grid size is loaded by the framework, the synapses and their annotations are automatically split into the respective pre- and postsynaptic sites, and the aggregate number of contact sites that share the same annotations is registered per subvolume.

### Null hypothesis for testing the impact of neuron morphology (step 2)

The parameter-free model of the null hypothesis (i.e., the null model *f*_*Null*_) computes synapse counts under the assumption that all combinations of connections between pre- and postsynaptic sites within a subvolume are equally likely. The expected synapse counts under the null model are *f*_*Null*_: λ_*a,b,k*_ = |a| ⋅ |b| ⋅ *n*^−1^ where |a| and |b| denote the number of pre- and postsynaptic sites for the respective grouping of neurons and/or subcellular compartments (e.g., |a| = |(INH, 812)| = 2 and |b| = |(EXC, 9123, D)| = 3) and *n*_*k*_ is the total synapse count in the subvolume *k*, i.e., ∑_a,b_ λ_*a,b,k*_ = *n*_*k*_.

### Synaptic specificity models and parameter inference (step 3)

A model with synaptic specificity *f*_α_ computes the expected number of synapses λ_*a,b,k*_ between groups of pre- and postsynaptic contact sites a, b in a subvolume: λ_*a,b,k*_(α) ∶= *f*_α_(λ^∗^). Models can be concatenated such that one model operates on the predicted number of synapses λ^∗^ from a previous model. Starting from the null model *f*_*Null*_, we defined a chain of models that represents increasing levels of synaptic specificity: *f*_*Null*_ → *f*_α*P*_ → *f*_α*PS*_ → *f*_α*PSC*_. Here, presynaptic sites have the attributes population *P* (e.g., *P* = {*EXC, INH*}) and/or cellular identity *C* = {i*d*_1_, i*d*_2_, … }. Postsynaptic sites can have an additional subcellular target site attribute *S* = {*S*, *D*, *A*} where S denotes soma, D dendrite, and A the AIS. The possible groupings of presynaptic sites a and postsynaptic sites b are the respective Cartesian products of these attributes, i.e., a ∈ *P* × *C* and b ∈ *P* × *C* × *S*. Specificity models at population level *f*_α*P*_ can introduce specificity parameters α_*P*_ for example by cell type, layer location, etc. Here, we test population level specificity by grouping the neurons according to their cell types as EXC and INH neurons (i.e., *P* = {*EXC, INH*}). Thus, every pairing results in 4 specificity parameters: α_*P*_ = (α_*EXC*,*EXC*_, α_*EXC*,*INH*_, α_*INH*,*EXC*_, α_*INH*,*INH*_). These cell type specificity parameters α_*xy*_ adjust the respective synapse counts from the null model: *f*_α*P*_: λ_*a,b,k*_^null^ = α_*xy*_ ⋅ λ where a ∈ {×} × *C* and b ∈ {*y*} × *C* × *S*. Specificity models at subcellular level *f*_α*PS*_ can introduce additional parameters by subcellular properties, for example the subcellular target site on postsynaptic neurons as we did here: α_*S*_ = (α_*Soma*_, α_*Dend*_, α_*AIS*_). These combined cell type and subcellular specificity parameters adjust the synapse counts from the preceding model: *f*_α*PS*_: λ_*a,b,k*_ = α_×_ ⋅ λ where a ∈ *P* × *C* and b ∈ *P* × *C* × {×}. Specificity models at cellular levels *f*_α*PSC*_ can introduce additional parameters by cellular identity, for example correlations between pairs of neurons as we did here based on the identity of the postsynaptic neuron α_*C*_ = (α_i*d*1_, α_i*d*2_, …). These combined cell type, subcellular and cellular specificity parameters adjust the synapse counts from the preceding model: *f*_α*PSC*_ where a ∈ *P* × *C* and b ∈ *P* × {×} × *S*. The technical range of the specificity parameters α_i_ is [0, M] and maps to the negative/positive preference interval of [−1, +1] as follows: α^′^ ≔ α_i_ − 1 (if α_i_ < 1) and α^′^ ≔ (α_i_ − 1)/(M − 1), otherwise. Synaptic specificity models can have parameters that are not analytically tractable. Simulation-based Bayesian inference (SBI)^39–41^ allows inferring suitable parameters for analytically intractable models from a set of simulation runs with sampled parameters. The prerequisites for applying SBI are (1) a stochastic simulator (y|α) that takes parameters α as input and generates observable statistics y, (2) access to ground truth observed statistics *y*_0_, and (3) a prior distribution over model parameters *p*(α). SBI uses a set of parameters sampled from the prior and corresponding simulated data {α_i_, *y*_i_}^*N*^ to train an artificial neural network-based density estimator to approximate the posterior distribution. After training, the density estimator is applied to the observed data *y*_0_ to obtain the posterior distribution over model parameters *p*(α|*y*_0_). Importantly, the inferred posterior distribution gives access to all data-compatible parameters and their covariance structure, enabling to analyze parameter uncertainties and interactions^29, 42^. For introductory usage examples of the SBI toolkit^43^, we refer to <u>sbi-dev/sbi: Simulation-based inference toolkit</u>. We employ SBI to identify specificity parameters α_a,b_ where a, b are pre- and postsynaptic groupings of neurons based on their cell type and/or subcellular compartments, for example, α_*P*_ = [α_*EXC*,*EXC*_, α_*EXC*,*INH*_, α_*INH*,*EXC*_, α_*INH*,*INH*_]. Based on aggregate statistics of the empirically observed synapse counts *y*_0_ = [*n*_*EXC*,*EXC*_, *n*_*EXC*,*INH*_, *n*_*INH*,*EXC*_, *n*_*INH*,*INH*_], SBI yields the posterior distribution over parameters *p*(α_*P*_|*y*_0_). Within each subvolume, we can compute local specificity parameters α_*a,b,k*_ where *k* denotes the subvolume. This can be done analytically by solving λ^*E*^_*a,b,k*_ = α_*a,b,k*_ ⋅ λ^*prec*^ for α_*a,b,k*_ where λ^*E*^ is the empirical synapse count (for the grouping of pre- and postsynaptic sites a, b in subvolume *k*) and λ^*prec*^ the synapse count predicted by the preceding model in the chain of models. The inferred specificity parameters α_*P*_ obtained by sampling from the posterior distribution *p*(α_*P*_|*y*_0_) were in good agreement with the average of the locally optimal parameters 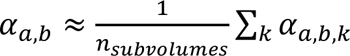 (**Fig. S5**).

### Quality metric for synaptic specificity models (step 4)

We define a loss function to quantify the predictive quality of a synaptic specificity model and the corresponding synaptic specificity parameters as follows. Let *n*_a,b_ be the empirically observed number of synapses from a presynaptic neuron a onto the specific compartment of a postsynaptic neuron b and let λ_a,b_ = ∑_*k*_ λ_*a,b,k*_ be the corresponding model-predicted synapse count summed over all subvolumes *k*. Assuming a Poisson random process, the probability that a connects with at least one synapse to b is *p*_a,b_ = 1 − exp(−λ_a,b_) and *p*^-^_a,b_ = 1 − *p*_a,b_. We define the model loss as:

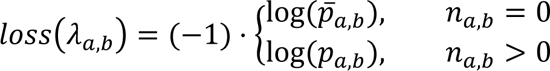

We measure the improvement of a specificity model α to the *Null* model by computing the delta loss, i.e., 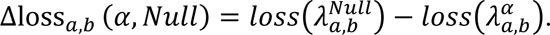

### Generation of discrete connectome realizations (step 5)

To generate connectome realizations from model-predicted synapse counts λ_*a,b,k*_in the subvolumes, we sample discrete synapse counts *n*_*a,b,k*_from a Poisson distribution (i.e., *n*_*a,b,k*_ ∼ *Poisson*(λ_*a,b,k*_)) unless λ_*a,b,k*_is itself discrete (i.e., when no local combinatorial variability is left). To compute the distributions of correctly predicted synapses, we generated 1000 connectome realizations for different specificity models. The histograms show how often observed synapses were predicted correctly when averaged over all realizations. The boxplots show the percentages of correctly predicted synapses per realized connectome. Aggregate connectivity statistics were computed based on discrete realizations of the model-generated connectomes. Aggregate synapse counts were directly obtained by grouping synapses according to the respective attributes (e.g., pre- and postsynaptic cell type). Synapse clusters refer to connections between annotated neurons (see step 1) in both datasets. We count all occurrences of the 16 triplet motifs between the selected neurons using a GPU-based implementation and compared these occurrences with those of an equivalent random network.

### Cell type clustering based on synaptic specificity

We restricted our analysis to neurons from the mouse dataset with at least 3 outgoing synapses to other annotated cells, resulting in n=127 neurons (109 excitatory, 18 inhibitory). We defined a specificity model that is agnostic to the known presynaptic cell type, thereby simulating a scenario in which these neurons are either unclassified or represent incompletely traced axon fragments. Based on the synaptic specificity parameters determined by the framework, we defined a *X* ≔ *n* × 5 feature matrix in which the rows represent neurons and the columns their specificity values onto: (1) EXC cells, (2) INH cells, (3) soma, (4) dendrite, and (5) AIS. Using scikit-learn we performed a standard scaling of the features and generated the dendrograms with the linkage function from scipy (Ward method with optimal leaf ordering). The PCA-embedding was performed with sklearn. After computing the dendrogram and embedding, the known cell type labels from the dataset were added post hoc for reference.

### Comparative analysis in human temporal cortex

Our analysis is based on the C3 stage segmentation of the human dataset^14^ with 133.7 million synapses. This segmentations comprises incompletely traced neurites, especially axons^44^. Connectivity between neurons at the cellular level is therefore underestimated in the currently available human dataset. To account for this limitation, we restricted our analysis to 200 neurons in layer 3 which were synaptically connected. For comparison with the mouse dataset, we refer to cells that are labeled ‘pyramidal’ as EXC and to cells labeled ‘interneuron’ as INH. We partitioned the volume innervated by the dendrites and axons of these 200 neurons into 9184 subvolumes using a 25 μm grid, which containing 36.8 of the 133.7 million synapses. We then performed the respective steps of the synaptic specificity analysis as previously described for the mouse dataset.

### Visualization and statistical analysis

Bar and line plots show the median and the 25th and 75th percentiles. The colored specificity matrices show the mean values of the respective synaptic specificity parameter distributions, mapped to the diverging blue/red color scale with the TwoSlopeNorm function from matplotlib. All charts were generated with matplotlib in combination with the numpy and sciPy libraries. Anatomical renderings of cells from the mouse and human dataset were generated with Amira^45^.

## Data and Code availability

All data is deposited to zenodo <u>10.5281/zenodo.14507538</u>. Our method is available as a computational framework, with source code and documentation via https://github.com/zibneuro/connectomedf. This repository comprises also a demo version of the code to reproduce examples shown in the figures. The mouse^27^ and human^14^ datasets are available at https://doi.org/10.5281/zenodo.3710458 and http://h01-release.storage.googleapis.com/landing.html, respectively.

## Acknowledgements

Funding was provided by the Max Planck Institute for Neurobiology of Behavior (MO), and by grants from the European Research Council: 633428, 101069192 (MO) and 101089288 (JM), the Deutsche Forschungsgemeinschaft: SFB 1089 (MO) and SPP 2041 (MO, JM, DB, HH), the German Federal Ministry of Education and Research: 01GQ1002, 01IS18052 (MO) and 01IS18039A (JM), a MLCoE grant 390727645 (JM), and a Neuroscience Network NRW grant iBehave (MO). We declare that we have no competing interests.

## Supplementary Figures and Figure Legends

**Figure S1:**
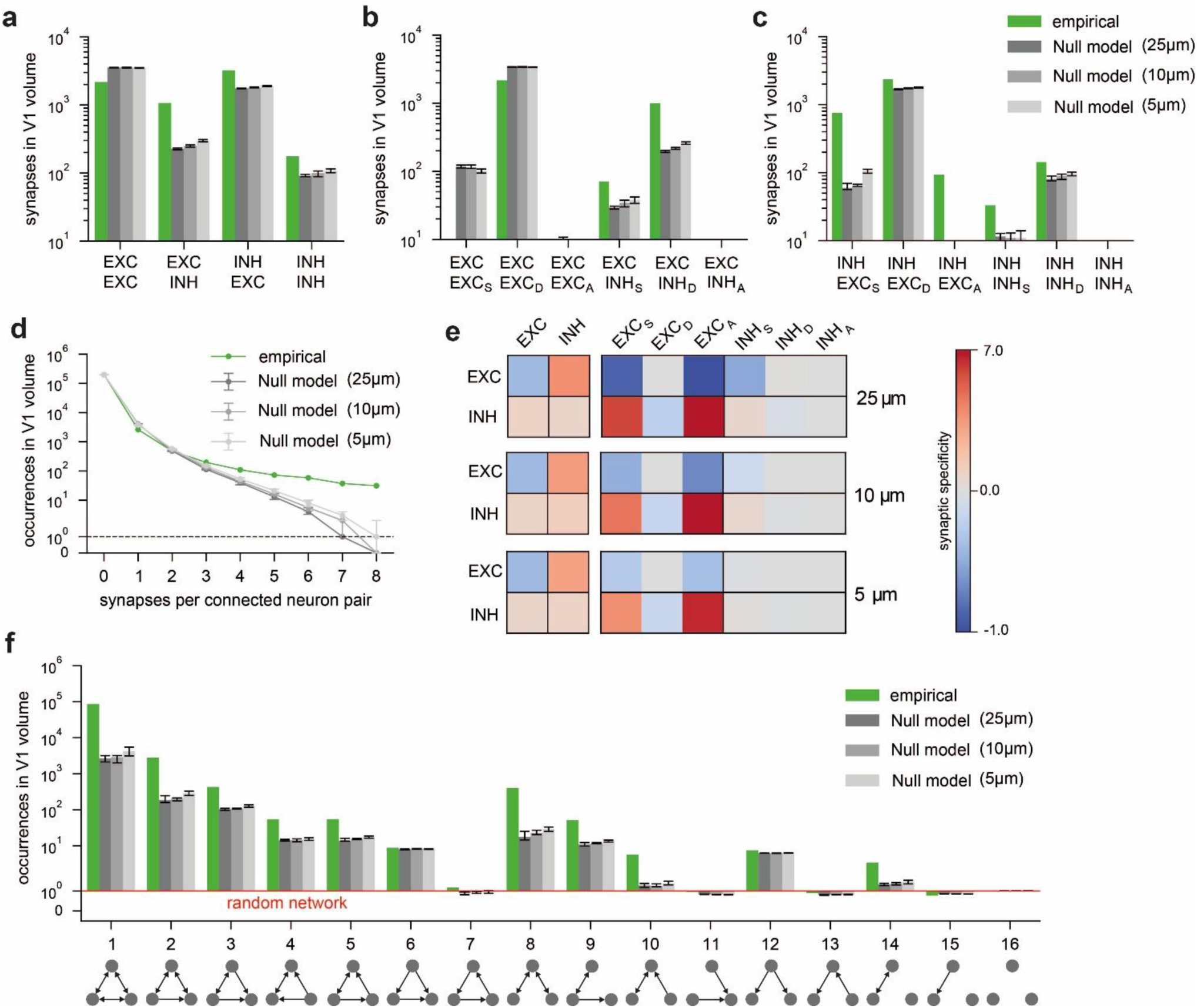
Testing the impact of neuron morphology on wiring specificity in mouse V1 with different grid sizes. **(a-f)** We repeated our null hypothesis analyses that we did for a grid size of 5 µm cubes in Fig. 3 for grid sizes of 10 and 25 µm cubes. The impact of neuron morphology on connectivity, and the inferred synaptic specify values – i.e., α parameters that model wiring conditional on neuronal properties – were remarkably robust with respect to these grid sizes (see also ^1^). **(a)** Synapse counts between EXC and INH neuron populations. **(b)** Synapse counts from presynaptic excitatory neurons disaggregated by postsynaptic cell type and compartment (S: soma, D: dendrite, A: axon initial segment). **(c)** Synapse counts from inhibitory neurons disaggregated by postsynaptic cell type and compartment. **(d)** Number of synapses per connected neuron pair. **(e)** Resulting synaptic specificity parameters when determined with different subvolume sizes. **(f)** Triplet motif occurrences.

**Figure S2:**
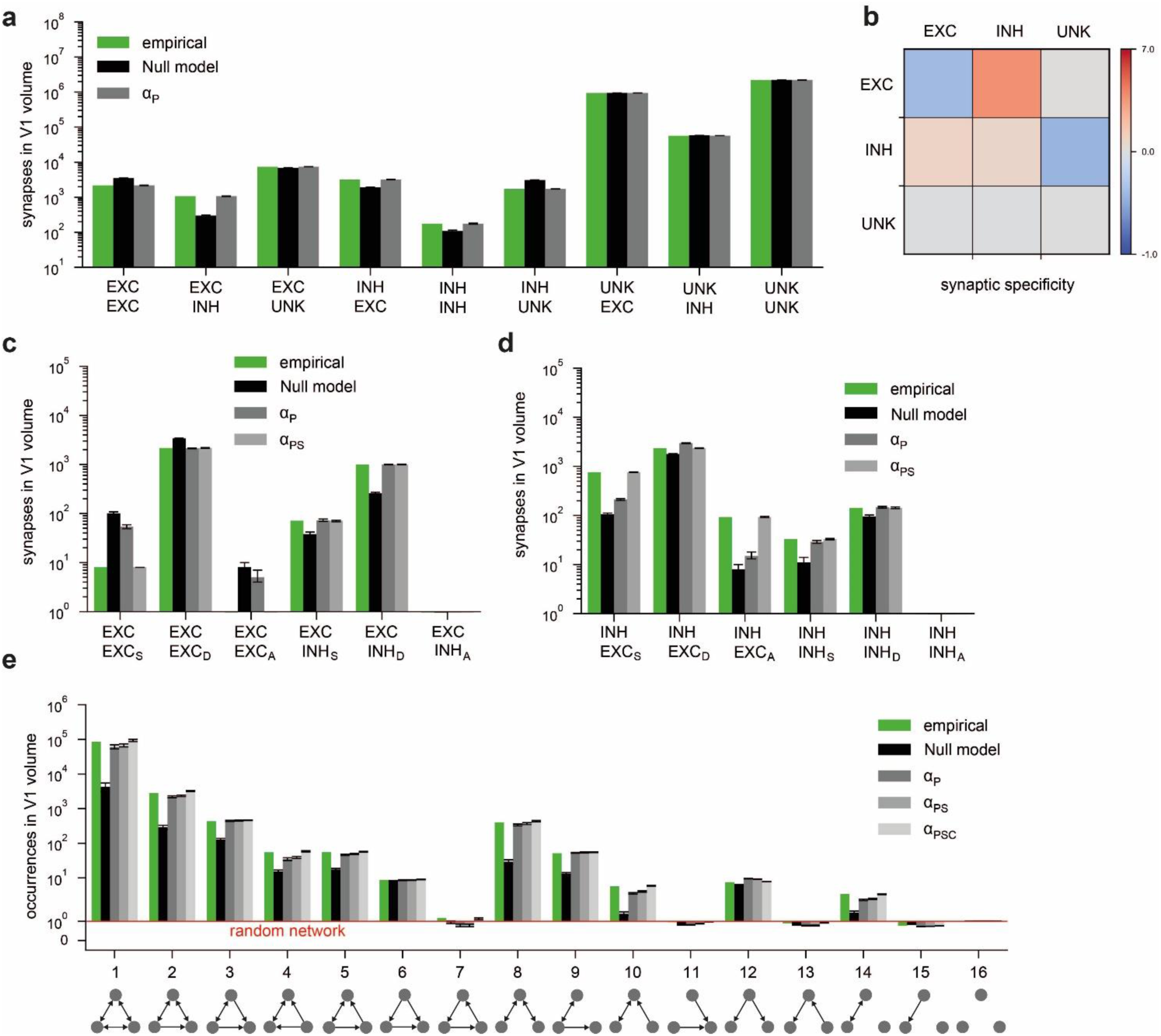
Testing the impact of synaptic specificity on wiring specificity in mouse V1. **(a-e)** Extension of the analyses in Fig. 4. **(a)** Synapse counts between neuron populations (EXC: excitatory, INH: inhibitory, UNK: unknown – i.e., representing synapses on neuronal structures that are not connected to a soma that is located within the reconstructed volume). **(b)** Synaptic specificity parameters based on the α_*P*_ model. **(c)** Synapse counts from presynaptic excitatory neurons disaggregated by the subcellular compartment of the postsynaptic neuron (S: soma, D: dendrite, A: axon initial segment). **(d)** Synapse counts from presynaptic inhibitory neurons disaggregated by the postsynaptic compartment. **(e)** Occurrences of triplet motifs comparing empirically observed frequencies to the predictions of the *Null*, α_*P*_, α_*PS*_, α_*PSC*_ models.

**Figure S3:**
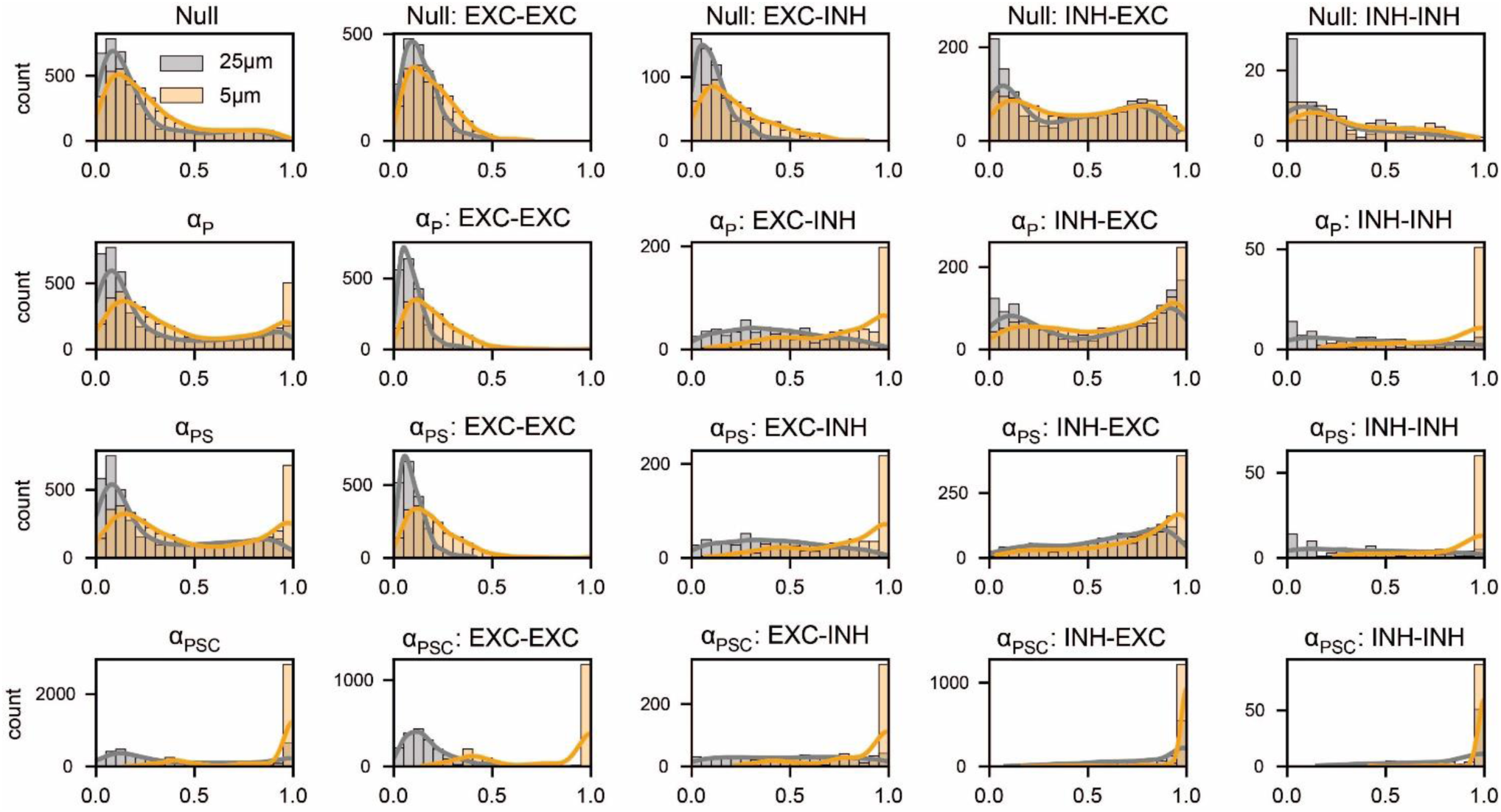
Testing the impact of synaptic specificity on the connectome from mouse V1. The distributions show the fractions of correctly predicted neuron-to-neuron connections (i.e., a neuron pair connected in the ground truth connectome is also connected in the model) in n=1000 realizations of the connectome under the given model. The distributions in the leftmost columns represent all neuron pairs; in the four columns on the right, the neuron pairs are separated by cell type. Model complexity increases by row (*Null*, α_*P*_, α_*PS*_, α_*PSC*_) and, as a result, the masses of the distributions move to the right (i.e., towards higher accuracy). **Please note:** We did this analysis for grid sizes of 5 (yellow) and 25 µm cubes (grey). While the impact of neuron morphology on connectivity, and inferred synaptic specify values were largely independent of grid size **(Fig. S1)**, predictability of empirically observed connectomes improves with smaller grid size.

**Figure S4:**
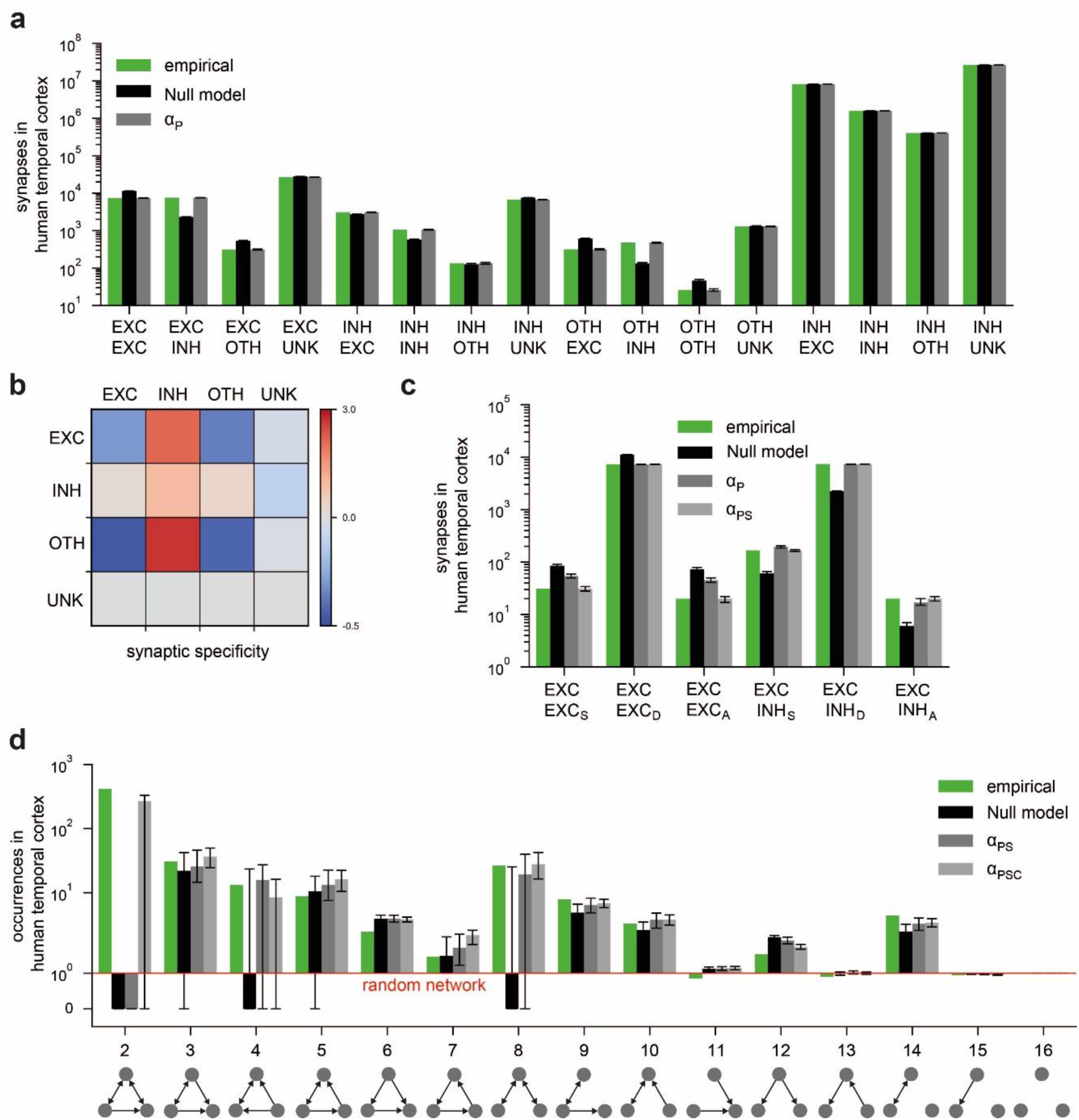
Testing the impact of synaptic specificity on wiring specificity in the human temporal cortex. **(a-e)** Extension of the analyses in Fig. 8. **(a)** Synapse counts between neuron populations grouped by cell type (EXC: excitatory, INH: inhibitory, OTH: other – i.e., neuronal structures that are not annotated as EXC or INH in the dataset, UNK: unknown – i.e., representing synapses on neuronal structures that are not connected to a soma that is located within the reconstructed volume). **(b)** Synaptic specificity values based on the α_*P*_model. **(c)** Synapse counts from presynaptic excitatory neurons disaggregated by postsynaptic cell type and subcellular compartment (S: soma, D: dendrite, A: axon initial segment). **(d)** Triplet motif occurrences among 200 neurons from layer 3.

**Figure S5:**
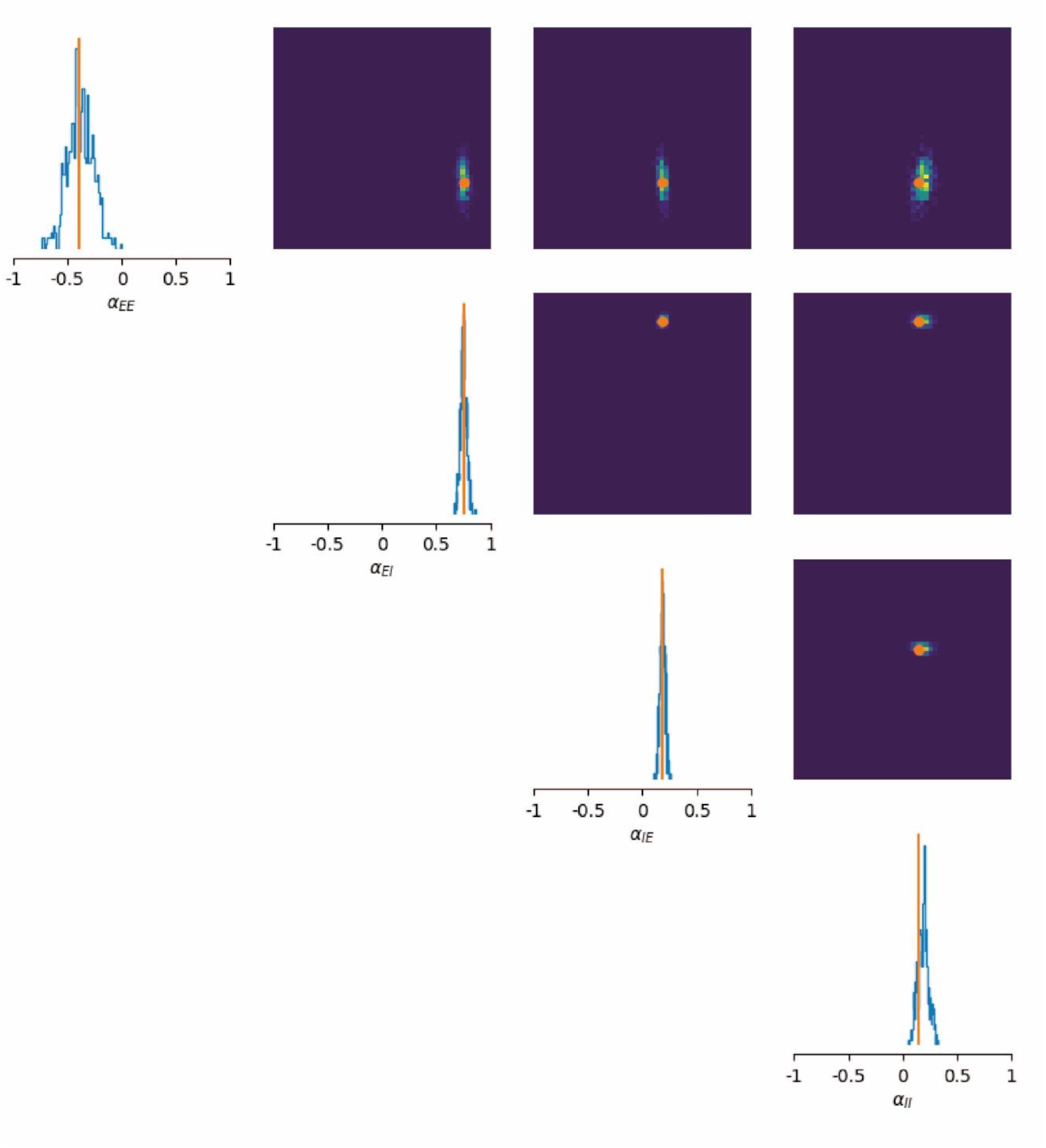
Simulation-based Bayesian parameter inference. Posterior distribution of inferred specificity parameters α_*P*_ = [α_*EE*_, α_*EI*_, α_*IE*_, α_*II*_]. Orange markers denote average values of analytically determined specificity parameters per subvolume (see Methods and ^2^).

